# Different computations underlie overt and covert spatial attention

**DOI:** 10.1101/2020.07.22.215905

**Authors:** Hsin-Hung Li, Jasmine Pan, Marisa Carrasco

**Author notes:** Correspondence: Hsin-Hung Li.

## Abstract

Perception and action are tightly coupled: visual responses at the saccade target are enhanced right before saccade onset. This phenomenon, presaccadic attention, is a form of overt attention—deployment of visual attention with concurrent eye movements. Presaccadic attention is well-documented, but its underlying computational process remains unknown. This is in stark contrast with covert attention—deployment of visual attention without concurrent eye movements—for which the computational process is well characterized. Here, a series of psychophysical experiments reveal that presaccadic attention modulates visual performance only via response gain changes even when attention field size increases, violating the predictions of a normalization model of attention, which has been widely used to explain the computations underlying covert attention. Our empirical results and model comparisons reveal that the perceptual modulations by overt and covert spatial attention are mediated through different computations.

## INTRODUCTION

Humans and primates make large and rapid eye movements, saccades, multiple times per second to explore visual scenes. Already before saccade onset, visual performance is improved (e.g., *d*′ in discrimination tasks) (e.g., Hoffman and Subramaniam, 1995; Kowler et al., 1995; Deubel and Schneider, 1996; Montagnini and Castet, 2007; Deubel, 2008; Collins et al., 2010; Rolfs and Carrasco, 2012; Hanning et al., 2019) and featural representations are modulated (Li et al., 2016; Ohl et al., 2017; Li et al., 2019) at the saccade target location. These effects, due to presaccadic attention, reveal a strong coupling between perception and action.

Presaccadic attention is a type of *overt attention*, or the deployment of visual attention with concurrent eye movements. Even though behavioral (Deubel and Schneider, 1996; Deubel, 2008; Rolfs and Carrasco, 2012; Li et al., 2016) and neural correlates (Fischer and Boch, 1981; Moore et al., 1998; Mazer and Gallant, 2003; Steinmetz and Moore, 2014; Moore and Zirnsak, 2017) of presaccadic attention have been extensively investigated, the computational process underlying the modulations of presaccadic attention remains unknown. This is in stark contrast with the research of *covert attention*, the deployment of attention without concurrent eye movements, for which not only the behavioral (Carrasco, 2011; Carrasco and Barbot, 2015) and neural correlates (Anton-Erxleben and Carrasco, 2013; Maunsell, 2015; Moore and Zirnsak, 2017) have been extensively investigated, but also the underlying computations (e.g., Boynton, 2009; Lee and Maunsell, 2009; Reynolds and Heeger, 2009; Kanashiro et al., 2017; Verhoef and Maunsell, 2017). To address this knowledge gap, here we ask: Can a unified computational framework account for the perceptual modulations by both overt presaccadic and covert spatial attention?

Characterizing how attention modulates the visual system’s input-and-output functions –in the present study specifically defined as visual performance as a function of stimulus contrast– is an essential step toward understanding the computations underlying attentional modulations. Contrast response functions have often been used to investigate attentional modulations. Several neurophysiological (McAdams and Maunsell, 1999; Reynolds et al., 2000; Martinez-Trujillo and Treue, 2002; Williford and Maunsell, 2006; Sani et al., 2017), psychophysical (Lu and Dosher, 1998; Dosher and Lu, 2000; Cameron et al., 2002; Morrone et al., 2004; Huang and Dobkins, 2005; Ling and Carrasco, 2006; Pestilli et al., 2007; Pestilli et al., 2009; Herrmann et al., 2010; Herrmann et al., 2012), and human neuroimaging (Buracas and Boynton, 2007; Li et al., 2008; Murray, 2008; Lu et al., 2011; Pestilli et al., 2011) studies have investigated how covert spatial attention and feature-based attention affect contrast response functions. Overall, the contrast response functions measured under various forms of covert attention can be explained by the Reynolds-Heeger Normalization Model of Attention (NMA) (Reynolds and Heeger, 2009) (also see, Boynton, 2009; Lee and Maunsell, 2009; Schwedhelm et al., 2016; Ni and Maunsell, 2017).

In this NMA, the attentional modulation is modeled as attentional gain factors that increase or decrease the input drive of visual neurons. The modulation by the attentional gain factors precedes normalization, and thus this modulation affects both the excitatory inputs and the suppressive inputs (the normalization pool) of visual neurons. This principle has been applied to explain attentional modulations in many neurophysiological studies (Reynolds and Heeger, 2009; Ni et al., 2012; Verhoef and Maunsell, 2016; Ni and Maunsell, 2019).

This NMA made several predictions regarding how covert attention modulates contrast response functions, which have been confirmed by empirical psychophysical findings: Spatial covert attention can modulate neural responses by horizontally shifting the contrast response function, changing the threshold of the psychometric function (contrast gain: quantified as a changes of semi-saturation contrast; **Figure 1A**). Spatial covert attention can increase neural responses by a multiplicative gain factor, improving behavioral performance by vertically scaling the psychometric function (response gain: quantified by a change of asymptotic response; **Figure 1B**). NMA and empirical studies have shown that spatial covert attention can induce a response gain change, a contrast gain change or a mix of both, depending on the size of the attention field relative to the size of the stimulus (**Figure 1**) (Reynolds and Heeger, 2009; Herrmann et al., 2010).

**Figure 1.**
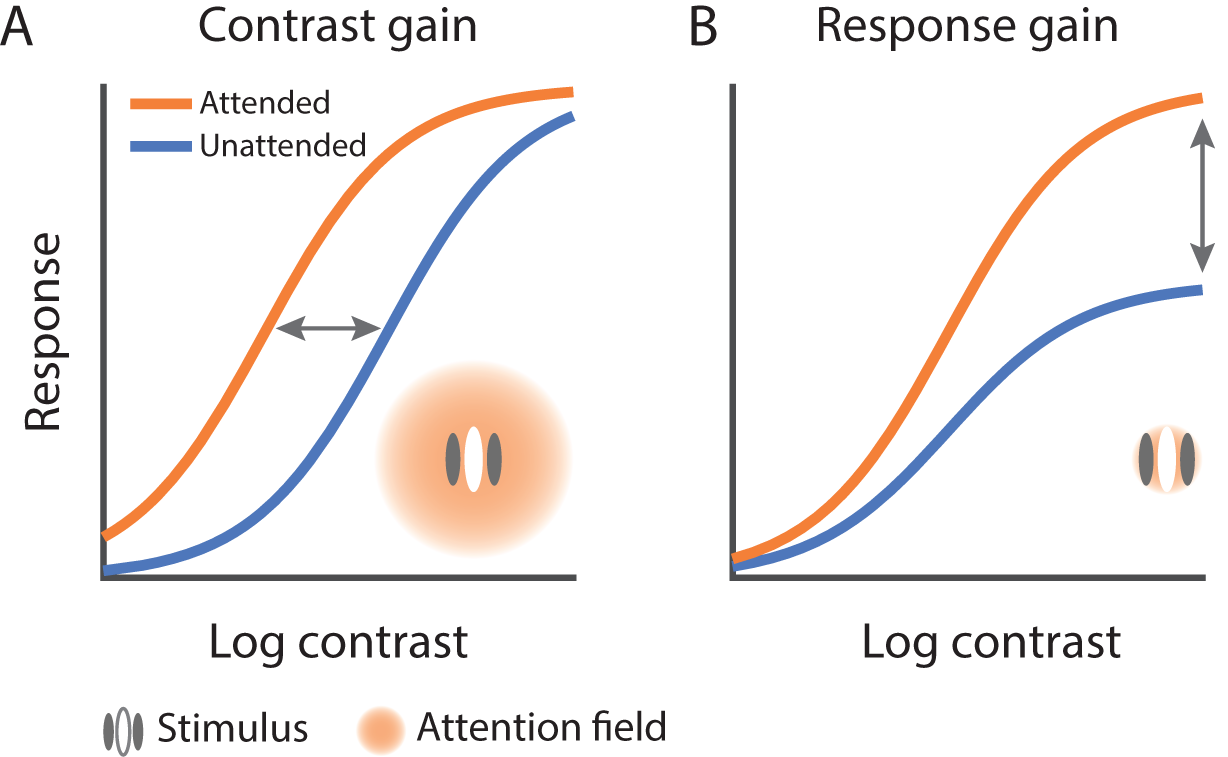
Different forms of attentional modulation on contrast response functions. (**A**) Contrast gain change: Attention horizontally shifts the response function as if attention scales the input contrast. (B) Response gain change: Attention scales neural response by a multiplicative gain factor, modulating the asymptotic response of the neuron. The normalization model of attention (NMA, Reynolds and Heeger, 2009) predicts that attention exhibits different modulations depending on the relative size of the attention field and the stimulus. A large attention field size (relative to the stimulus size) leads to contrast gain changes (**A**); a small attention field size leads to response gain changes (**B**)

To elucidate the computation underlying presaccadic attention, here we characterize the effect of presaccadic attention on the contrast response function in a series of psychophysical experiments. In Experiment 1, we investigated the effect of presaccadic attention when only one stimulus, positioned at the saccade target location, was presented. In Experiment 2, we assessed the effect of presaccadic attention at both the saccade target and non-target locations, and we compared presaccadic attention with the effect of covert endogenous and covert exogenous attention under the same stimulus parameters. In Experiment 3, we investigated whether the effect of presaccadic attention depends on the location uncertainty of the target stimulus. In all cases, the effects of presaccadic attention were compared to a neutral baseline in which no saccadic eye movements were executed.

To preview our results, we found that presaccadic attention only generated a response gain change, whereas covert attention generated a mix of response gain and contrast gain changes. The response gain change generated by presaccadic attention was robust even when we increased the location uncertainty of the target stimulus. By fitting various computational models to our data, we found that a model in which attention modulates the neural response by a gain factor multiplicatively after normalization outperformed the NMA. Conversely, the NMA outperformed other models in explaining the perceptual modulations by covert attention. These results reveal that the gain modulations by overt presaccadic attention are mediated through neural computations that are different from those previously proposed and empirically confirmed for covert attention.

## RESULTS

### Experiment 1

We characterized the modulation of presaccadic attention on the contrast response function when only one stimulus was presented at the saccade target location. Observers were asked to discriminate the orientation of the target, presented 8° left or right of fixation. To map out the psychometric function, the contrast of the target varied trial-by-trial. In the neutral condition, a neutral pre-cue instructed observers to maintain fixation at the screen center throughout the trial. In the saccade condition, a saccadic pre-cue instructed observers to saccade to the upcoming target location (**Figure 2A**), in which the target was presented shortly after the onset of the cue. Observers had to report the target orientation, left or right of vertical. Gaze positions were monitored throughout the trials.

**Figure 2.**
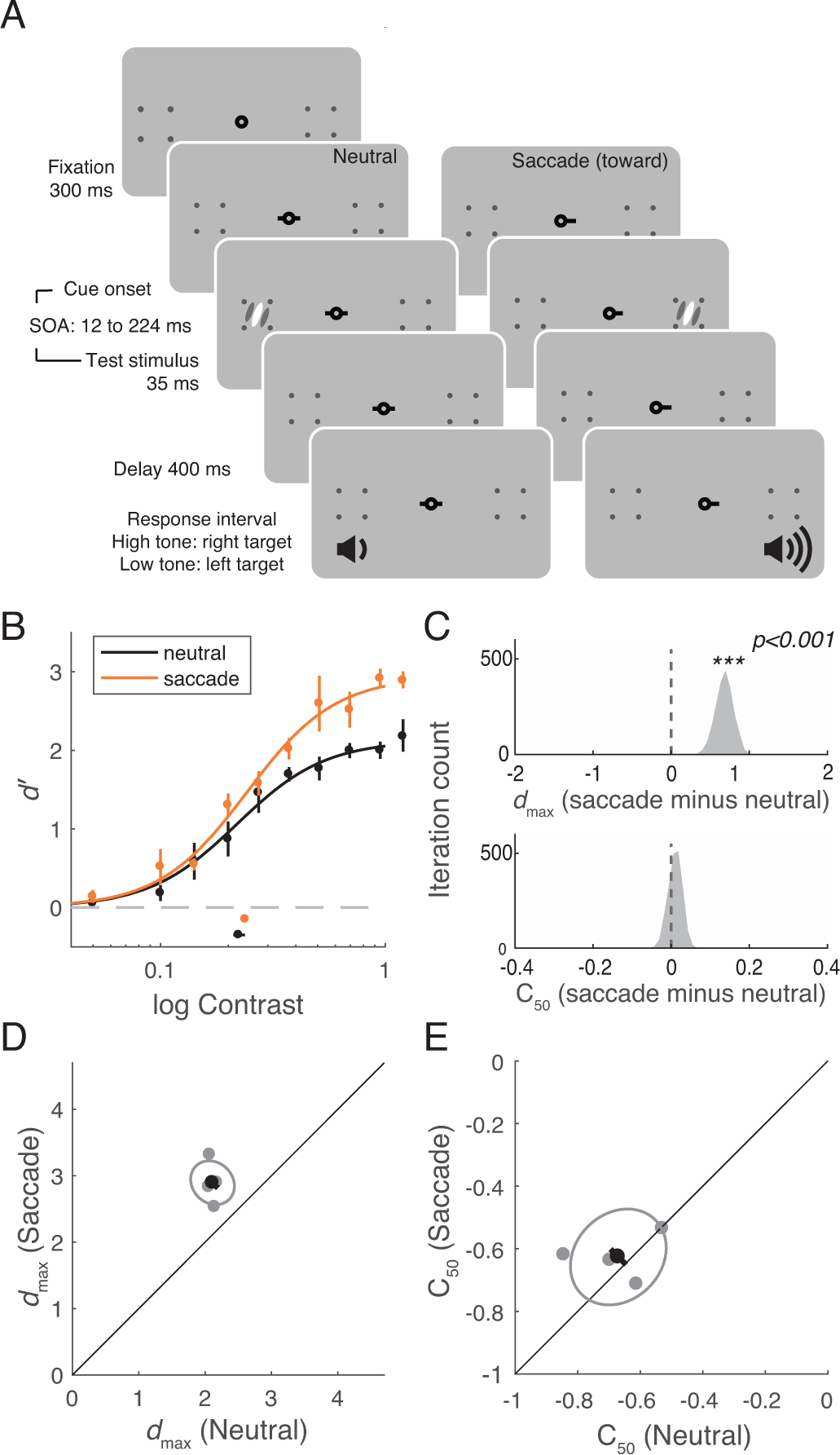
Experiment 1. (**A**) Procedures. Each trial started with a fixation period. A neutral cue or a saccadic cue directed observers to either maintain fixation or make a saccade toward the upcoming target location. The target appeared shortly after the cue onset. Observer had to report the orientation of the target after hearing an auditory response cue. Two tones in different frequencies were used to inform the observers the location of the target. (**B**) Group-averaged psychometric functions (*d*′vs. contrast). *d*_*max*_ and *C*_*50*_ of the group-averaged psychometric functions are plotted at the right and the bottom of the figure. The error bars represent 95% bootstrapped confidence interval. (**C**) The bootstrapped distribution of the difference of *d*_*max*_ (top) and the difference of C50 (bottom) between the neutral and saccade conditions. Asterisks denote that the distribution is significantly different from zero. (**D**) Gray dots: Best-fitted *d*_*max*_ of individual observers; Gray ellipse: the ellipse with major axes oriented toward the difference and the sum of the *d*_*max*_ across two conditions. The difference (or sum) of *d*_*max*_ is first computed for individual observers, and the major axis of the ellipse represent ±1 standard deviation of the difference (or sum); The black dot: group-averaged *d*_*max*_ and the error bar represents the standard deviation of the bootstrapped distribution of the difference between neutral and saccade conditions. (**E**) C50 illustrated in the same format as in (D).

Trials in the saccade condition were sorted offline based on the relative timing between saccade onset and stimulus offset (**Supplementary Figure 1**). We compared the performance in the neutral condition to the performance in a presaccadic time window (75 ms to 0 ms before saccade onset; Rolfs and Carrasco, 2012; Szinte et al., 2015; Li et al., 2016; Li et al., 2019). We fitted performance (*d′*) as a function of target contrast with Naka-Rushton functions (see **Methods**). Presaccadic attention enhanced performance at the saccade target through an increment of *d*_*max*_, the asymptotic level of performance (*p*<0.001, bootstrapping test). *C*_*50*_, the semi-saturation constant, was not significantly different between the two conditions (*p*=0.81, bootstrapping test) (**Figure 2B-2E**). These results are consistent with a response gain change.

### Experiment 2

We investigated the effect of presaccadic attention at both the saccade-target location and a non-target location horizontally opposite to the saccade direction. In addition, we compared the effects of presaccadic attention to the effects of covert spatial attention. Observers performed three different tasks in separate sessions: A presaccadic attention, a covert endogenous (voluntary) attention and a covert exogenous (involuntary) attention tasks. We compared the three types of attention under the same stimulus parameters, while concurrently allowing each type of attention to take full effect by using their optimal cue location and cue timing (see **Methods** and **Figure 3**).

**Figure 3.**
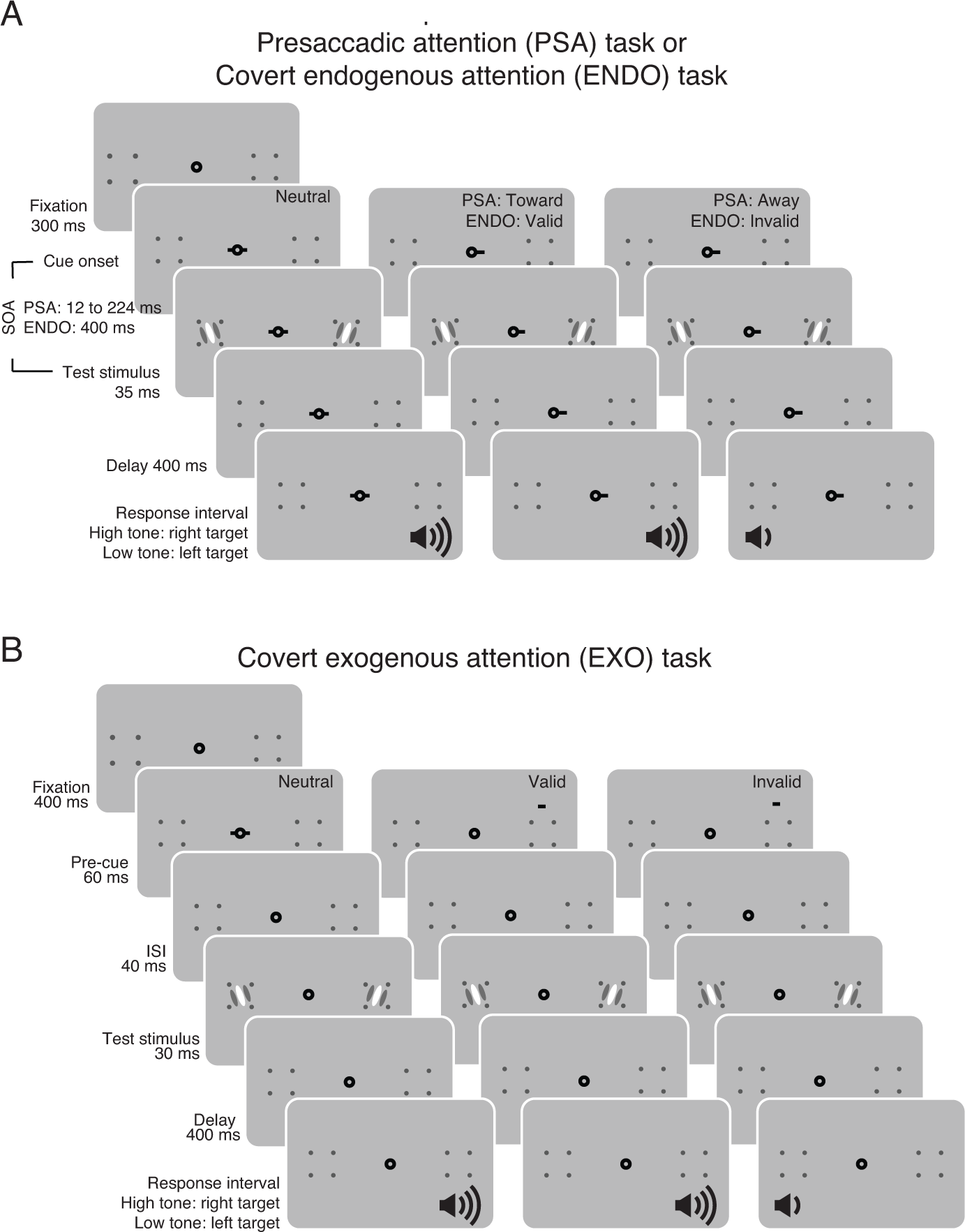
Experiment 2. Observers were tested with three tasks in different sessions: Presaccadic attention (PSA) task, Covert endogenous attention (ENDO) task and Covert exogenous attention (EXO) task. (**A**) Here we illustrate the procedures of the PSA task and the ENDO task together as they shared a similar stimulus sequence. The cue-to-target SOA was shorter for the PSA task than in the ENDO task. Two stimuli, a target and a distractor, were presented simultaneously. In the PSA task, the pre-cue instructed observers to saccade to the cued location. In the saccade Toward condition the pre-cue matched the target location whereas in the saccade Away condition the pre-cue indicated the distractor. Saccade Toward and saccade Away conditions were interleaved with equal probability and thus the saccade cue was uninformative regarding the target location. In the ENDO task, observers maintained fixation throughout the trials and the pre-cue instructed observers to covertly pay attention to the cued location. The valid and invalid trials were interleaved within blocks with 70% and 30% probability respectively; thus, the pre-cue was informative regarding the target location. At the end of a trial, observers reported the orientation of the target probed by the auditory response cue (high tone: target at the right; low tone: target at the left). (**B**) In the EXO task, the pre-cue was a brief flash. In the neutral condition, the pre-cue was presented at the fixation. In the valid condition, the pre-cue was flashed above the target whereas in the invalid condition, the pre-cue was presented above the distractor. The valid and the invalid trials were interleaved with equal probability, and thus the pre-cue was uninformative regarding the target location. See details in Methods.

For the presaccadic attention task, two stimuli (a target and a distractor), one in the left and one in the right visual field, were presented simultaneously in each trial (**Figure 3A**). In the saccade condition, a saccadic pre-cue instructed observers to saccade either toward the target (the Toward condition) or away from the target, to the distractor (the Away condition) with equal probability. Observers had to report the orientation of the target indicated by an auditory response cue at stimulus offset. This design allowed us to investigate the effects of presaccadic attention without conflating saccade direction with cue validity as the saccadic pre-cue did not convey any information regarding the relevance of the stimuli.

In the endogenous attention task, covert attention was manipulated by an informative (70% validity) central pre-cue. The cue indicated the target location in the valid trials and indicated the distractor in the invalid trials (**Figure 3A**). In the exogenous attention task, an uninformative peripheral pre-cue was flashed adjacent to the target or the distractor with equal probability (**Figure 3B**). Observers maintained fixation at the screen center throughout each trial. Only in the endogenous attention task, observers were instructed to pay attention to the cued location.

We observed both a benefit and a cost of presaccadic attention. Compared to the neutral condition, when averaging *d′* across contrast levels, performance (*d*′) was improved in the Toward condition (*p*<0.001, bootstrapping test) and was impaired in the Away condition (*p*<0.001, bootstrapping test). Summarizing the effect of presaccadic attention by comparing the fit of the Naka-Rushton functions between the Toward and the Away conditions, we observed a significant change of *d*_*max*_ (*p*<0.005, bootstrapping test) without a significant change of *C*_*50*_ (*p*=0.15, bootstrapping test) (**Figure 4A-4D**).

**Figure 4.**
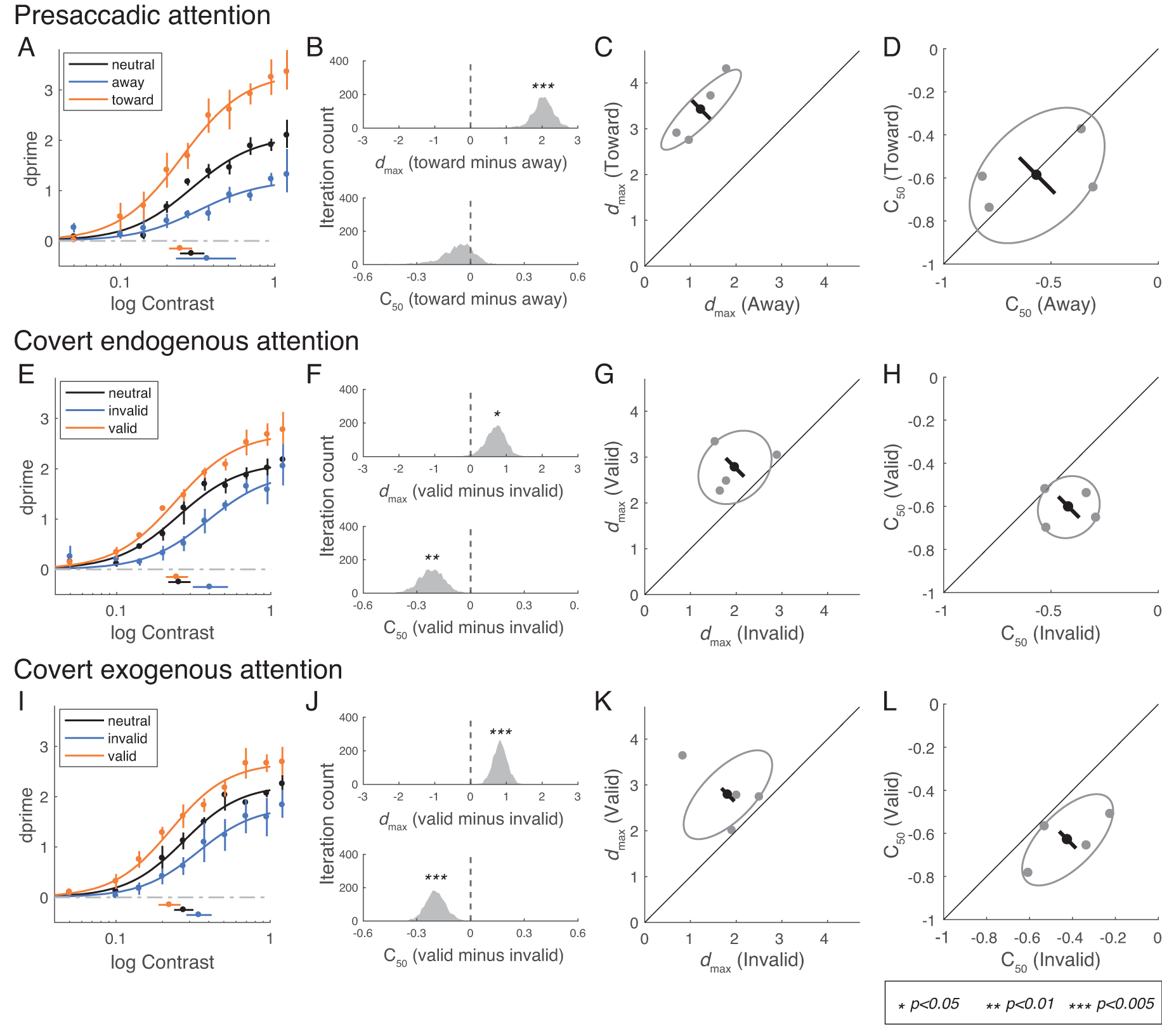
Experiment 2. (**A-D**) Saccade condition. (**A**) Group-averaged psychometric functions (d′ vs. contrast). *d*_*max*_ and *C*_*50*_ of the group-averaged psychometric functions are plotted to the right and bottom of the figure. The error bars represent the 95% bootstrapped confidence interval. (**B**) The bootstrapped distribution of the difference of *d*_*max*_ (top) and the difference of C50 (bottom) between the Toward and Away conditions. Asterisks denote that the distribution is significantly different from zero. (**C**) Gray dots: Best-fitted *d*_*max*_ of individual observers; Gray ellipse: the ellipse with major axes oriented toward the difference and the sum of the *d*_*max*_ across two conditions. The difference (or sum) of *d*_*max*_ is first computed for individual observers, and the major axis of the ellipse represent ±1 standard deviation of the difference (or sum); The black dot: group-averaged *d*_*max*_ and the error bar represents the standard deviation of the bootstrapped distribution of the difference between Toward and Away conditions. (**D**) *C*_*50*_ illustrated in the same format as in (C). (**E-H**) Covert endogenous attention condition; corresponding to A-D. (**I-L**) Covert exogenous attention condition; corresponding to A-D and E-H.

Covert endogenous attention also led to an improvement of performance in the valid condition (*p*<0.001, bootstrapping test) and a decrement of performance in the invalid condition (*p*<0.001, bootstrapping test). Different from the effect of presaccadic attention, when comparing the fit of Naka-Rushton functions between the valid and the invalid conditions, we observed changes in both *d*_*max*_ (*p*<0.05, bootstrapping test) and *C*_*50*_ (*p*<0.01, bootstrapping test; **Figure 4E-4H**).

The effects of covert exogenous attention condition were similar to those of covert endogenous attention. Compared to the neutral condition, performance was higher in the valid condition (*p*<0.001, bootstrapping test) and lower in the invalid condition (*p*<0.001, bootstrapping test) than the neutral condition. Both *d*_*max*_ and *C*_*50*_ showed significant changes when comparing the valid and invalid conditions (both *p*<0.001, bootstrapping test) (**Figure 4I-4L**).

In sum, we found that overt and covert attention modulated performance through different gain changes. Covert attention generated a mix of response gain and contrast gain changes, whereas presaccadic attention only generated a response gain change.

### Experiment 3

Both the NMA (Reynolds and Heeger, 2009) and empirical results (Herrmann et al., 2010; Herrmann et al., 2012) have demonstrated that the modulations generated by covert attention (both exogenous and endogenous) are affected by the size of the attention field relative to the stimulus. When the attention field size is large compared to the stimulus, attention generates a contrast gain change, and when the attention field size is small, attention generates a response gain change. Thus, the findings that presaccadic attention generated a pure response gain change in Experiment 2 might reflect that presaccadic attention was deployed locally at the saccade target location whereas the deployment of covert spatial attention was not as precise, implicating a larger attention field size compared to presaccadic attention.

Were the effect of presaccadic attention to follow the predictions of NMA, contrast gain changes would emerge with an enlarged attentional field size. To test this prediction, following the protocols in covert attention studies (Herrmann et al., 2010; Cutrone et al., 2018), we increased stimulus location uncertainty. This manipulation encourages observers to pay attention to a larger spatial area. It is possible that sometimes we may be preparing a saccade to a particular location, but we may want to monitor a larger area in case the object of interest is around, but not exactly at, the saccade location.

We considered the presaccadic attention task in Experiment 2 as a low-uncertainty condition. In Experiment 3, we measured the effect of presaccadic attention in a medium-uncertainty and a high-uncertainty conditions, in which the target (and the distractor) was randomly presented in one of the pre-designated locations in each trial (**Figure 5A and 5B**). We asked observers to still saccade to the center of the placeholder. Following NMA, the effect of presaccadic attention should transform from a pure response gain change (in Experiment 1 and 2) to a contrast gain change (or a mix of both) in the medium- and high-uncertainty condition.

**Figure 5.**
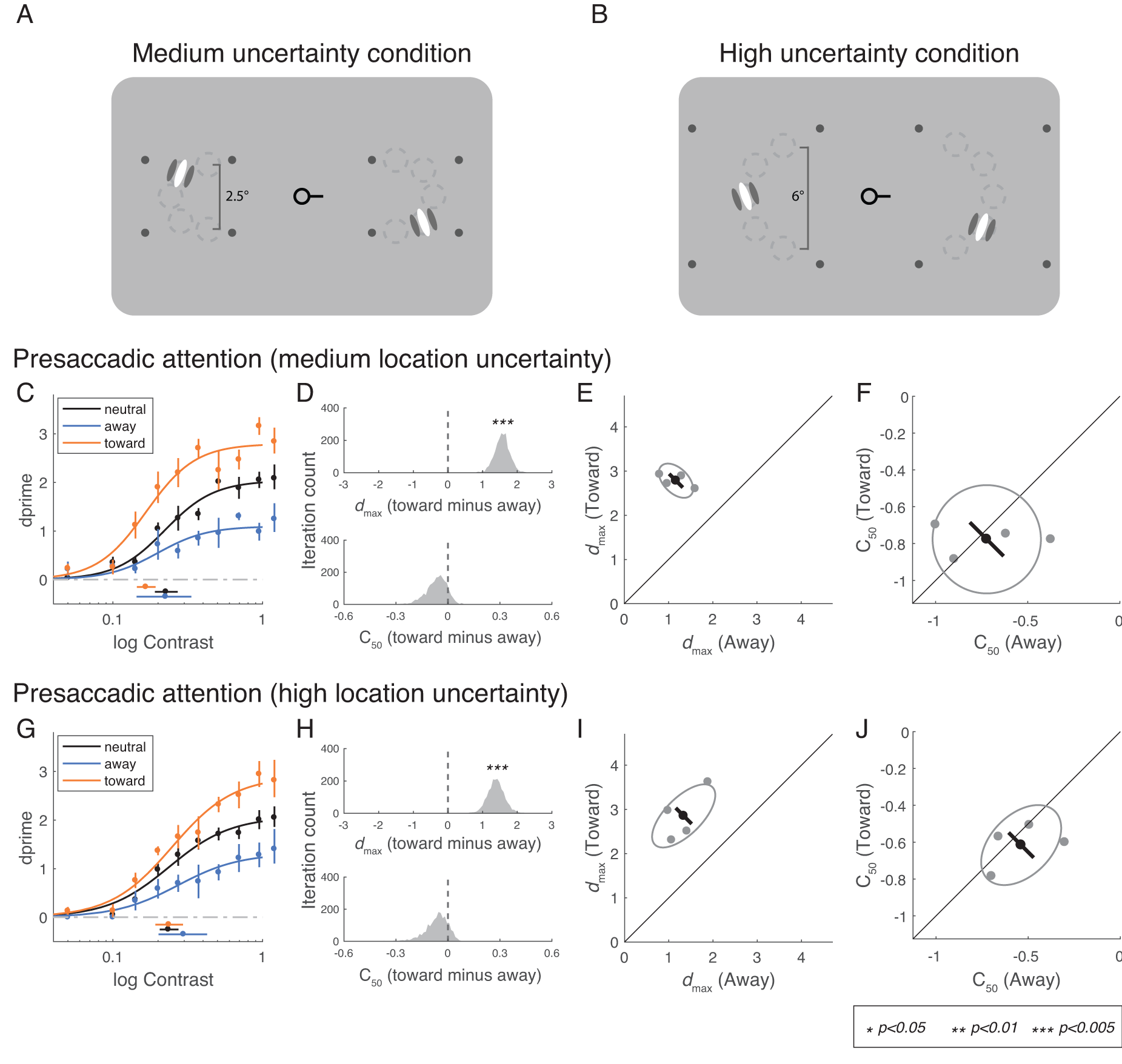
Experiment 3. **(A-B)** The procedure of Experiment 3 is the same as the presaccadic attention task in Experiment 2, except that the test stimuli were presented with location uncertainty. Stimuli appeared at one of five predefined iso-eccentric locations. Stimulus locations were randomly and independently selected in the left and right placeholder. The dashed circles (not displayed during the experiments) indicate possible stimulus locations (five in each placeholder; the figures here are for the purpose of illustration only. See Methods for the details of the stimulus parameters). **(C-F)** Medium-uncertainty condition. (**C**) Group-averaged psychometric functions (d′ vs. contrast). *d*_*max*_ and *C*_*50*_ of the group-averaged psychometric functions are plotted at the right and bottom of the figure. The error bars represent 95% bootstrapped confidence interval. (**D**) The bootstrapped distribution of the difference of *d*_*max*_ (top) and the difference of *C*_*50*_ (bottom) between the Toward and Away conditions. The distribution significant different from zero is denoted with asterisks. (**E**) Gray dots: Best-fitted *d*_*max*_ of individual observers; Gray ellipse: the ellipse with major axes oriented toward the difference and the sum of the *d*_*max*_ across two conditions. The difference (or sum) of *d*_*max*_ is first computed for individual observers, and the major axis of the ellipse represent ±1 standard deviation of the difference (or sum); The black dot: group-averaged *d*_*max*_ and the error bar represents the standard deviation of the bootstrapped distribution of the difference between Toward and Away conditions. (**F**) C50 illustrated in the same format as in (**E**). **(G-J)** High-uncertainty condition, corresponding to C-F.

Presaccadic attention exhibited response gain changes even when location uncertainty increased. In the medium-uncertainty condition, *d*_*max*_ differed significantly between the Toward and the Away conditions (*p*<0.001, bootstrapping test) but the corresponding *C*_*50*_ did not (*p*=0.15, bootstrapping test) (**Figure 5C-5F**). In the high-uncertainty condition, the results were the same: there were significant changes in *d*_*max*_ (*p*<0.001, bootstrapping test), but not in *C*_*50*_ (*p*=0.19, bootstrapping test), between the Toward and the Away conditions (**Figure 5G-5J**).

We conducted simulations to compare the predictions of NMA with the data in Experiments 2 and 3. Simulations are based on Reynolds and Heeger’s NMA (2009) and the stimulus parameters of our experiments. To simulate the effect of location uncertainty, we scaled the width of attention field size from 0.5, 2.5 to 6 degrees to match the stimuli in the low-, medium- and high-uncertainty conditions respectively (**Supplementary Figure 2**). The model predicted that attentional modulations on neural responses shift from a response gain change to a contrast gain change with increasing attention field size (indexed by location uncertainty) (**Supplementary Figure 2**). Note that the simulation here is only to ensure that given the assumption of NMA, our stimuli, and the attention field size that increases with the location uncertainty, we should observe a shift from response gain to contrast gain changes, which was not observed in our data. We later conducted model comparisons to formally compare NMA and alternative models: response gain, contrast gain, baseline input and output baseline models.

### Pre-saccadic attention field size

Is it possible that observers’ attention field size did not increase with location uncertainty as we expected? For example, observers’ attention field might have remained small and localized at the center of the placeholder. Were this the case, the mismatch between the predictions of NMA and the empirical results would be due to the ineffectiveness of our experimental manipulations.

To investigate this possibility, we binned the trials in the high-uncertainty condition based on the target location: the two outmost locations, two intermediate locations and the central location (see legends in **Figure 6A, 6E and 6I**). We found significant changes in *d*_*max*_ between the Toward and the Away conditions at all target locations (outmost locations *p*<0.01; intermediate locations *p*<0.05; central location *p*<0.005, bootstrapping test). We did not observe significant *C*_*50*_ shifts at any of the locations (outmost locations *p*=0.61; intermediate locations *p*=0.10; central location *p*=0.22, bootstrapping test). In addition, there was no significant difference in the magnitude of response gain changes (quantified as the *d*_*max*_ in the Toward condition minus the *d*_*max*_ in the Away condition) across the three bins (central vs. outmost *p*=0.63; central vs. intermediate *p*=0.26; outmost vs. intermediate *p*=0.20, bootstrapping test). Moreover, the distributions of saccade landing points were almost constant regardless of the location at which the target was presented (**Supplementary Figure 3**). These results support the assumptions in the simulations that in the high-uncertainty condition, the size of attention field extends beyond the central target location.

**Figure 6.**
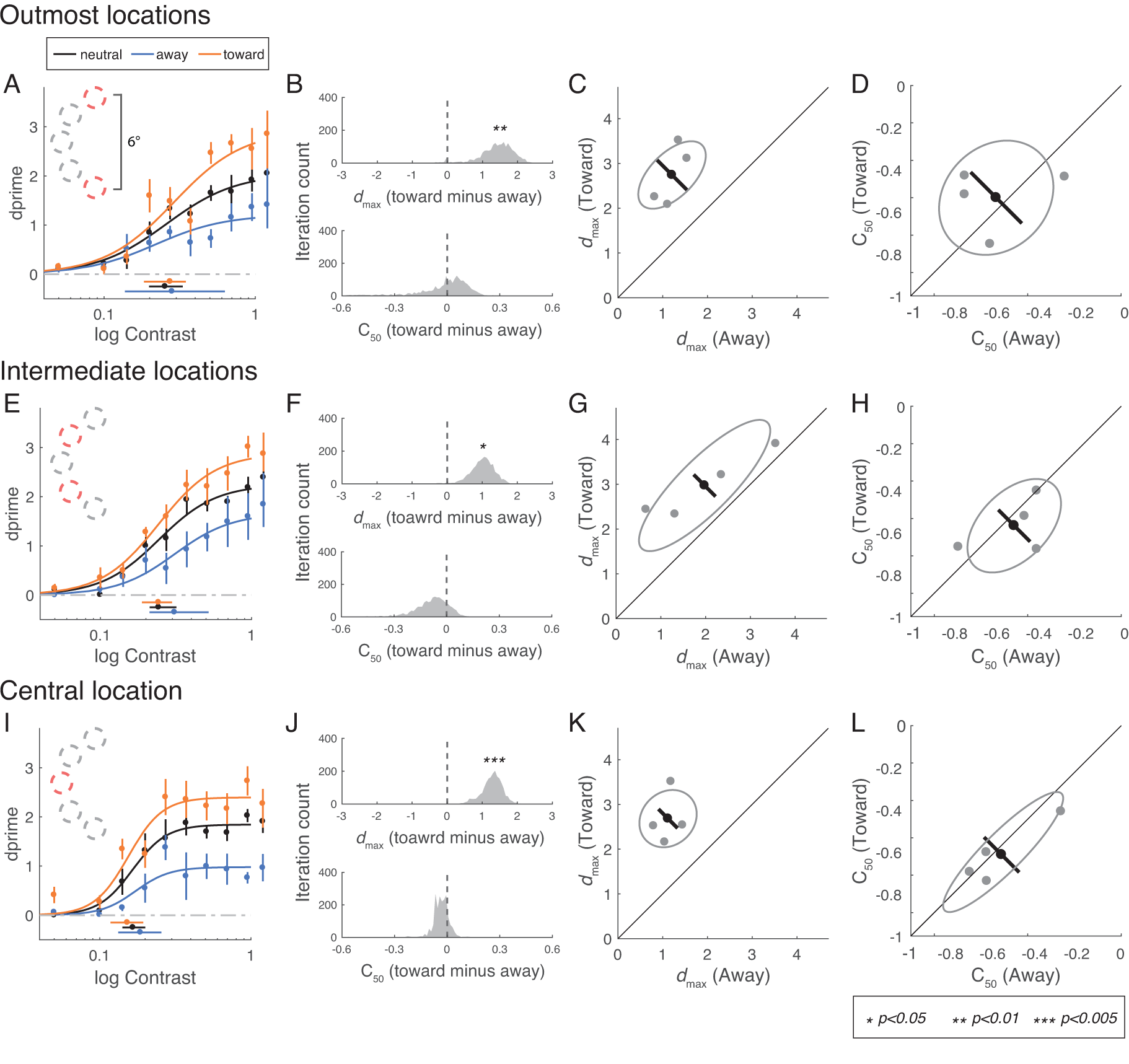
Performance binned by target location in high-uncertainty condition. **(A-D)** Outmost locations. (**A**) Group-averaged psychometric functions (d′ vs. contrast). *d*_*max*_ and *C*_*50*_ of the group-averaged psychometric functions are plotted at the right and the bottom of the figure. The error bars represent 95% bootstrapped confidence interval. (**B**) The bootstrapped distribution of the difference of *d*_*max*_ (top) and the difference of C50 (bottom) between the Toward and Away conditions. The distribution significant different from zero is denoted with asterisks. (**C**) Gray dots: Best-fitted *d*_*max*_ of individual observers; Gray ellipse: the ellipse with major axes oriented toward the difference and the sum of the *d*_*max*_ across two conditions. The difference (or sum) of *d*_*max*_ is first computed for individual observers, and the major axis of the ellipse represent ±1 standard deviation of the difference (or sum); The black dot: group-averaged *d*_*max*_ and the error bar represents the standard deviation of the bootstrapped distribution of the difference between Toward and Away conditions. (**D**) C50 illustrated in the same format as in (**C**). **(E-H)** Intermediate locations, corresponding to (**A-D**). **(E-H)** Central location, corresponding to (**A-D**).

### Model comparisons

The absence of size-dependent gain change by presaccadic attention in our experiments indicates that the modulations by presaccadic attention may not be mediated by the NMA (Reynolds and Heeger, 2009; Herrmann et al., 2010; Herrmann et al., 2012). We built and compared various computational models to explain the computations underlying the modulations of presaccadic attention. In the models, we computed population neural response and used signal detection theory to link neural response to behavioral reports. Doing so enabled us to estimate the likelihood of the models given the observer’s response on a trial-by-trial basis.

The models are based on the framework that neural response can be estimated by a normalization equation (Carandini and Heeger, 2012). In different models, attention modulates neural response through different parts or stages of normalization: In the response gain model, attention is modeled by response gain factors that multiplicatively modulate the response of the neurons after normalization (**Figure 7A**); In the NMA model, attention affects neural responses through attentional gain factors that multiplicatively modulate the excitatory drives of the neurons before normalization (**Figure 7C**) (Reynolds and Heeger, 2009); In the contrast gain model, attention divisively modulates the suppression constant in normalization (**Supplementary Figure 4**); In the input baseline model (Cutrone et al., 2014), attention modulates the excitatory inputs of the neurons additively prior to normalization (**Supplementary Figure 5**); In the output baseline model (Li et al., 2008; Hara et al., 2014), attention is modeled as an additive term after normalization (**Supplementary Figure 6**) (see details of all the models in **Methods**).

**Figure 7.**
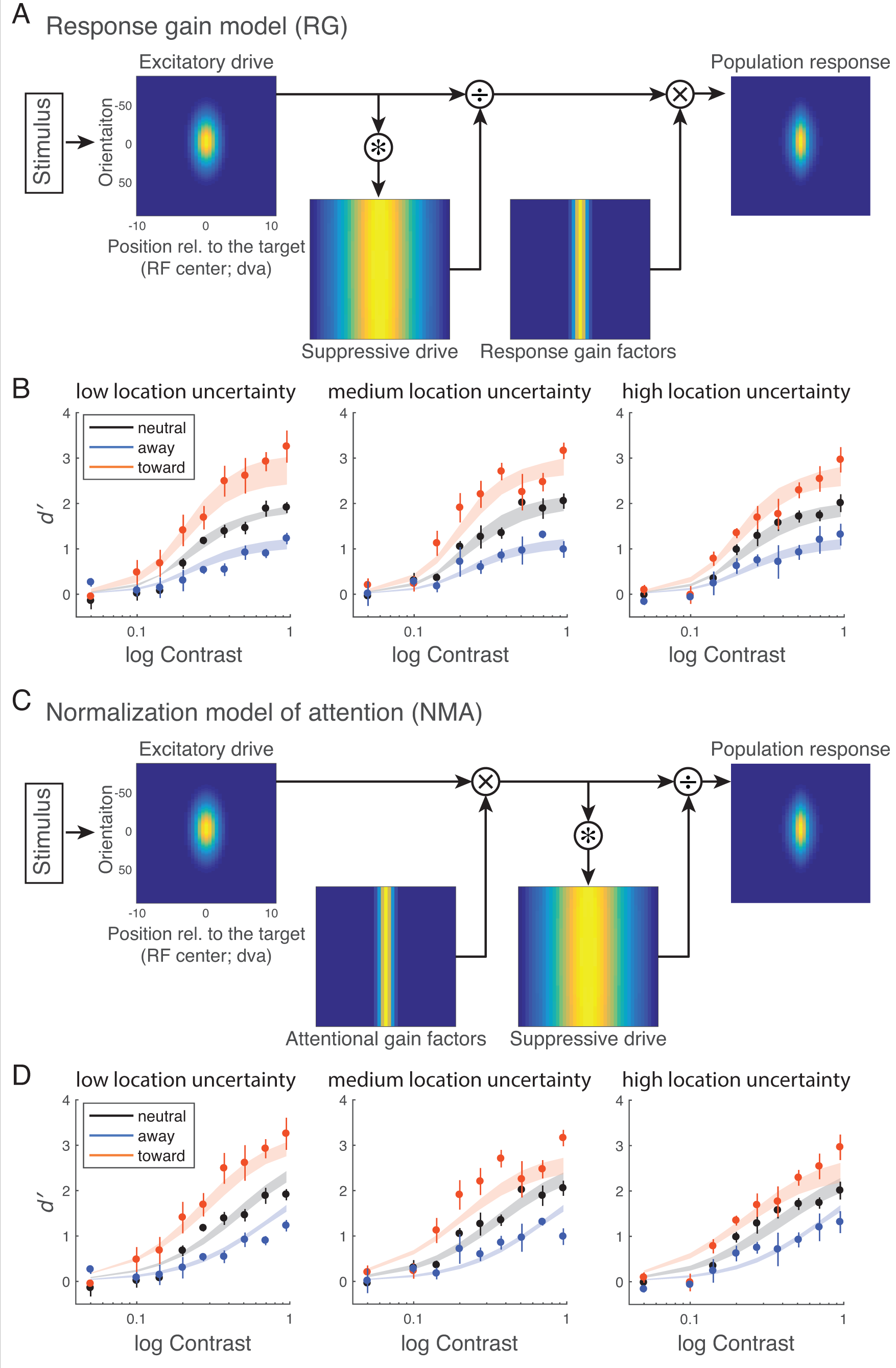
Models and models fits. (A) Response gain (RG) model. Attention multiplicatively modulates neural response after normalization. (B) Model fit of the RG model. Data points and error bars represent group-averaged data and ±1 s.e.m. The model was fitted to individual subjects, and the shading areas represent the average of the fits across participants (mean ±1 s.e.m). (C) Normalization model of attention (NMA). Attentional modulation based on Reynolds and Heeger’s normalization model of attention. Attention is modeled by attentional gain factors that modulate excitatory drive of the neurons before normalization. (D) Model fit of the NMA model. Data points and error bars represent group-averaged data and ±1 s.e.m. The model was fitted to individual subjects, and the shading areas represent the average of the fits across participants (mean ±1 s.e.m).

We jointly fitted the models to the data of the presaccadic attention condition across three levels of location uncertainty (low, Experiment 2; **Figure 4A**; medium, Experiment 2, **Figure 5C;** high, Experiment 2; **Figure 5G**). We fitted the free parameters to the data of each individual subject using maximum-likelihood estimation, and used Akaike Information Criterion (AIC; Akaike, 1974; Akaike, 1998) as the index for model comparison.

We found that the response gain model outperformed the NMA in fitting the data by a group-summed ΔAIC score of 84 (95% boostrapped CI [28, 216]) (**Figure 8**), whereas the output baseline model performed the worst. We conducted a factorial model comparison and tested 20 models in total. The ranking of the models was not specific to the setup of the other factors in the model: The response gain model was the best-fit model even when we changed the setup of the response bias or the trade-off between attention field size and the magnitude of attentional modulation (**Supplementary Figure 7**).

**Figure 8.**
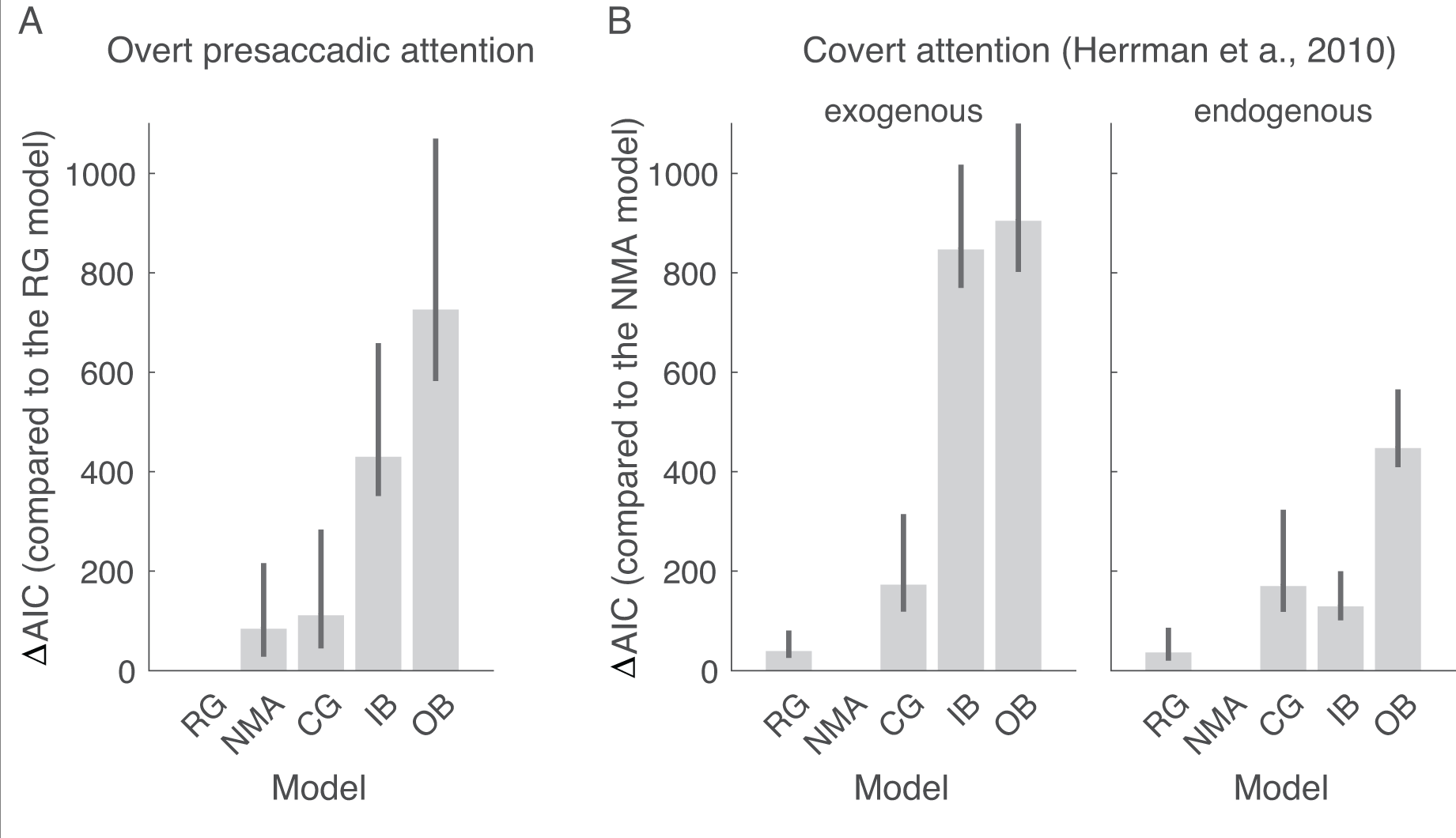
Model comparison using AIC. (A) Model comparisons for the presaccadic attention experiments. ΔAIC is the AIC of each model minus the AIC of the best-fit model (response gain, RG, model). The closer the ΔAIC is to zero, the better the model fits the data. The bars represent ΔAIC summed across participants. The error bars represent 95% bootstrapped confidence interval. (B) Model comparisons for the data reported in Herrmann et al., 2010. ΔAIC is the AIC of each model minus the AIC of the best-fit model (Normalization model of attention, NMA, model). RG: response gain; CG: contrast gain; IB: Input baseline; OB: output baseline. Here, all the models contain a response bias term *c* as a free parameter, without a trade-off term *p* (see details in **Methods** and **Supplementary Figure 7-9** for complete illustrations of the results of model comparisons).

We also fitted the models to the data of a previous study reporting the size-dependent gain changes induced by covert exogenous and covert endogenous attention (Herrmann et al., 2010). Herrmann et al. (2010) fitted Naka-Ruston functions to the psychometric functions (*d′* vs. contrast) without fitting process models like ours that incorporated both population neural response and behavioral reports, and generated predictions at a single-trial level. Critically, we found that different from the results of overt presaccadic attention, the NMA model is the best model to explain the effect of both exogenous and endogenous covert attention (ΔAIC between the NMA model and the RG model: endogenous attention: 36 [25, 82]; exogenous attention: 39 [25, 80]) (**Figure 8B**).

## DISCUSSION

We investigated the perceptual modulations of presaccadic attention, a form of overt attention, by characterizing its effect on performance as a function of stimulus contrast. We found that presaccadic attention exhibited a response gain change, whereas covert endogenous and covert exogenous attention exhibited a mixture of response gain and contrast gain change on the psychometric functions. The response gain change by presaccadic attention was robust even when location uncertainty of the stimuli increased. By model comparisons, we found that a model that assumes that overt presaccadic attention modulates neural responses after normalization—a response gain model—outperformed the Reynolds and Heeger’s NMA (Reynolds and Heeger, 2009), a computational model that has explained various phenomena of covert attention (Herrmann et al., 2010; Herrmann et al., 2012; Ni et al., 2012; Verhoef and Maunsell, 2016; Ni and Maunsell, 2019). These findings indicate that the computations underlying the modulations of these two types of spatial attention—overt presaccadic and covert—are different.

Previous studies have distinguished presaccadic attention and covert attention by creating experimental conditions that limit the effect of one type of attention but not the other. For example, presaccadic attention can take effect at a shorter latency than covert endogenous attention (Rolfs and Carrasco, 2012; Li et al., 2016), covert attention can facilitate performance for stimuli outside of the oculomotor range (Hanning et al., 2019), and observers can covertly attend to a task-relevant location even when a saccade was made to other locations (Klapetek et al., 2016). Here, while keeping experimental designs and stimulus parameters as similar as possible, we optimized cue location and timing to maximize the operation of each type of attention (**Figure 3**). Our results demonstrate that overt presaccadic and covert spatial attention not only differ in the conditions under which they can take effect, but also in the consequence of their deployment.

Previous psychophysical experiments of covert attention had found varying gain modulations across studies. For example, Huang and Dobkins (2005) reported that covert attention exhibited both contrast and response gain changes in contrast discrimination tasks; with dual-task procedure, Morrone et al. (2004) found response gain changes by attention for both luminance-defined and color-defined stimuli. Pestilli, Ling and Carrasco (2009) reported contrast gain for endogenous attention and response gain for exogenous attention. The NMA and later experiments testing its predictions (Herrmann et al., 2010, 2012) pointed out that the degree of response gain and contrast gain changes by covert attention depends on the attentional field size and the stimulus size (**Figure 1**). Thus, to obtain a pure response gain or contrast gain change for covert attention, one has to titrate the stimulus size and attentional field size in a specific manner. Our results that both endogenous and exogenous attention exhibited a mix of response gain and contrast gain change result from the stimulus parameters we used: The eccentricity at which the stimuli were presented, the size and eccentricity of the placeholders, and the size and eccentricity of the peripheral pre-cue for exogenous attention, all parameters which could influence the size of the attentional field, resulting in similar attention field sizes in both the exogenous and endogenous attention tasks.

Albeit the differences in their details (Reynolds and Heeger, 2009; Ni et al., 2012; Verhoef and Maunsell, 2016; Ni and Maunsell, 2019), in NMA, the computations of attention can be conceptualized by a normalization equation: *R* = *E*(*c*_*center*_) / [*S*(*c*_*center*_ + *c*_*surround*_) + *σ*], in which the visual neuron’s response *R* is determined by the excitatory drive *E*, suppressive drive (normalization pool) *S* and a nonnegative suppression constant *σ. E* is a function of the input contrast *c*_*center*_ at the location central to the neuron’s receptive field. Visual neurons have an extended suppressive surround and thus *S* is a function of the input contrast (*c*_*center*_ and *c*_*surround*_) at both the center and the surround. Critically in NMA, attention is modeled as a gain factor *a* that concurrently modulates excitatory and suppressive inputs prior to divisive normalization. When the attention field is large (covering both the center and the surround), neural response becomes *R* = *E*(*ac*_*center*_) / [*S*(*ac*_*center*_ + *ac*_*surround*_) + *σ*], leading to a contrast gain change as in **Figure 1A**; when attention field is small (without “attending to” the surround), the neural response is *R* = *E*(*ac*_*center*_) / [*S*(*ac*_*center*_ + *c*_*surround*_) + *σ*], leading to a response gain change as in **Figure 1B** (Reynolds and Heeger, 2009). The absence of size-dependent gain change by presaccadic attention in our experiments indicates that the modulations by presaccadic attention can instead be summarized by a response gain factor *γ* that scales neural responses after normalization *R* = *γ E*(*c*_*center*_) / [*S*(*c*_*center*_ + *c*_*surround*_) + *σ*].

The concepts above are confirmed by model comparisons. The model assuming that attention multiplicatively modulates neural response after normalization is best at explaining the effect of presaccadic overt attention. In contrast, we found that NMA is best in explaining the effect of both endogenous and exogenous covert attention. Thus, even though NMA has been used to explain many neural and perceptual modulations generated by covert attention our results suggest that such computation does not apply to presaccadic overt attention. A response gain mechanism could be a signature of the perceptual modulations rooted in saccade preparation.

The modeling approach we took is more comprehensive than the approach in the previous studies, in which the data were compared with the predictions of NMA by fitting Naka-Rushton functions to the psychometric functions (Herrmann et al., 2010; Herrmann et al., 2012). Instead of describing the data by parameter values *d*_*max*_ and *C*_*50*_, in our models we computed population neural response and made predictions on trial-by-trial behavioral reports. Among a neural population, depending on the neurons’ preferred orientation and location, different neurons could exhibit different magnitudes or forms of attentional modulations (Hara et al., 2014). This heterogeneity of the attentional effect was taken into account by the models because the signal available for the observers to make a decision encompasses the entire simulated neural population. In addition, this approach also allowed non-sensory or decision-related factors (e.g., response bias) to be fitted in the models.

Note that our models assumed that attentional field is maintained at the center of the placeholder regardless of location uncertainty. This assumption is in line with the findings that presaccadic attention depends on the planned location during oculomotor programming rather than the exact saccade landing point (Van der Stigchel and De Vries, 2015; Van der Stigchel and de Vries, 2018; Wollenberg et al., 2018). Moreover, in our data, the averaged saccade landing points changed very little (<0.13°) when the target was presented away from the central location (**Supplementary Figure 3**). Jittering the location of the simulated attention field by this amount led to negligible changes on the simulated neural responses.

It may be surprising that the NMA model did not fit the data of presaccadic attention well, as NMA could mimic either response gain or contrast gain by adjusting the size of the attention field to be either small or large, respectively. Note that our models take into account the stimulus location on a trial-by-trial basis. For the conditions with location uncertainty, if the size of the attention field remains small, there would be no attention effect at the outmost or intermediate locations, or attention could even generate a suppressive effect on the target, which was not observed in the data (**Figure 6**). The fits of NMA (**Figure 7D**) seem to indicate that the model used shallower (non-saturating) functions in an attempt to generate different maximum responses at the highest contrast level (a response-gain-like pattern) across the Toward, Away and neutral conditions. However, this led to model fits that cannot quite capture the shape of the psychometric function as well as the response gain model.

For NMA to explain our data, the following conditions would have had to be fulfilled: In the high-uncertainty condition, the attention field size would have to remain small, and attention would have to shift to the location wherever the target appeared during the presaccadic interval while participants were still able to saccade to the center of the placeholder. This is as if the target attracted stimulus-driven attention onto itself. This is unlikely for two reasons. First, this stimulus-driven attention would have to be extremely fast and only occur in the presence of presaccadic attention; otherwise the target in the endogenous and exogenous attention tasks would have had the same effect. Second, if the stimulus generated an attention-capture effect, saccade endpoints would have been affected, resulting in averaged saccades that landed between the planned saccade landing point and the target stimulus (Coren and Hoenig, 1972; Findlay, 1982). However, the distribution of saccade endpoints remained practically constant even when location uncertainty was high (**Supplementary Figure 3**).

The coupling between saccadic eye movements and visual attention are well-documented at both behavioral (Hoffman and Subramaniam, 1995; Kowler et al., 1995; Deubel and Schneider, 1996; Castet et al., 2006; Montagnini and Castet, 2007; Collins et al., 2010; Shimozaki et al., 2012; Hanning et al., 2018) and neural (Kustov and Robinson, 1996; Armstrong et al., 2006; Armstrong and Moore, 2007; Ikkai and Curtis, 2011; Zhao et al., 2012; Engel et al., 2016; Moore and Zirnsak, 2017) levels. Our results resonate with the notion that even though the neural correlates of saccade and covert attention are shared at a coarse scale, the two phenomena are neurophysiologically dissociable (Juan et al., 2004; Thompson et al., 2005; Gregoriou et al., 2012; Steinmetz and Moore, 2014; Lowe and Schall, 2018). For example, Gregoriou et al. (2012) reported that in the frontal-eye-field (FEF), visual and visuomotor cells showed enhanced firing rate during a covert attention task, whereas motor and visuomotor cells showed enhanced firing rate (time-locked to saccade onset) during a saccade task. In addition, the synchronization between FEF and visual cortex V4 also depends both on cell types and task. These distinct neural substrates underlying saccade execution and covert attention might contribute to the differential perceptual modulations and computations we observed for overt presaccadic attention and covert attention.

Our results are consistent with the findings that neural responses in sensory visual cortex are enhanced in a retinotopically specific manner (Fischer and Boch, 1981; Moore et al., 1998; Moore, 1999; Saber et al., 2015) prior to eye movements. This presaccadic enhancement is accompanied by an improvement of orientation decodability of saccade target (Moore and Chang, 2009). In a subsequent study Steinmetz and Moore (2014) reported that the enhancement of neural firing rate can occur at two locations simultaneously, a task-relevant location and a saccade-target location, indicating that attentional demand (task relevance) and presaccadic enhancement are dissociable. However, in this study the modulations by saccade preparation and attention (relevancy) are very similar in both their form (increased firing rate) and magnitude. This is probably not surprising as they aligned the neural response to cue onset, and analyzed an epoch long before (> 800 ms) saccade execution. Moreover, given our results, in order to reveal the differences between the modulations by presaccadic attention and covert attention, a finer characterization of the modulations is called for by mapping out the entire contrast response functions just before saccade onset. Future neurophysiological studies can take this approach to investigate whether the neural modulations by overt and covert attention in visual cortex are supported by different computations as our behavioral and computational results indicate.

To conclude, this study reveals for the first time that presaccadic attention and covert attention are mediated by different computations. Whereas a well-established model of covert attention, the NMA, does not account for the computations underlying the effects of presaccadic attention on contrast sensitivity, a response gain model does.

## Acknowledgments

NIH Grant R01EY019693, members of the Carrasco Lab, in particular Antonio Fernández, Nina Hanning and Michael Jigo.

## Methods

### Participants

Four observers (two females and two males. Age range: 22-33) participated in the three experiments (a total of 21 1-hr sessions per observer). All observers had normal or corrected-to-normal vision. The experiment was conducted with the written consent of each observer and the University Committee on Activities involving Human Subjects at New York University approved the experimental protocols.

### Setup

Observers sat in a dimly lit room with the chin rest positioned 57 cm from the monitor. Presentation of the stimuli was controlled by MATLAB (Mathworks) using the Psychophysics Toolbox extensions (Kleiner et al., 2007). The monitor was gamma-corrected and had a resolution of 1280 × 960 pixels with a refresh rate of 85 Hz. The gaze position of the right eye was recorded by an EyeLink 1000 with the Desktop Mount (SR Research).

### Experiment 1

Each trial started with a fixation period of 300 ms followed by the onset of a neutral cue in the neutral condition or a saccadic cue in the saccade condition. The saccadic cue was a bar at the fixation pointing toward the upcoming target location (8° left or right from the fixation; Figure **2A**). Two placeholders, marking the two potential target locations, were presented throughout the experiment. Observers were instructed to make a saccadic eye movement to the cued location as fast as possible when they saw the saccadic cue. The neutral cue was composed of two bars pointing to both locations, instructing observers to maintain fixation at the center throughout the trials. The test stimulus (the target, a Gabor presented for 35 ms) appeared shortly after the cue onset. Eye position was monitored throughout the trials.

The SOA (stimulus onset asynchrony) between the cue and the target was randomly sampled between 12 and 224 ms in each trial. This range of SOA was used because it allowed the test stimulus to be presented before saccade onset in most of the trials in the saccade condition. Moreover, this interval was shorter than the time (∼300 ms) required for covert endogenous spatial attention to take effect (Carrasco, 2011). After the offset of the test stimulus and a delay of 400 ms, an auditory tone was played as the response cue. Tones in two different frequencies informed the observer at which location (left or right) the target had been presented. After hearing the response cue, the observer reported the orientation of the target (left vs. right tilt from vertical) by pressing either the left-arrow or right-arrow key on the keyboard. Observers were given 3 s to respond after the response cue. After the button press, the next trial started after an inter-trial-interval ranging from 0.8 to 1.2 s. The neutral condition and the saccade condition were tested in separate blocks, in a randomized order. On average, every observer performed 2592 trials in three sessions on different days.

### Experiment 2

Three types of attention were manipulated in Experiment 2: Presaccadic attention, covert endogenous (voluntary) attention and covert exogenous (involuntary) attention. On average, every observer completed 8892 trials in 11 sessions on different days. The procedures for the presaccadic attention task were identical to the ones used in Experiment 1, except the following modifications: The test stimuli in Experiment 2 were composed of two Gabors, a target and a distractor, one presented 8° to the left and one presented 8° to the right of fixation simultaneously (**Figure 3A**). The location of the target was determined randomly in each trial. In the saccade condition, the saccadic cue either pointed toward the target (Toward condition) or the distractor (Away condition) with equal probability. Thus, the observer had to saccade to the distractor (i.e., away from the target) in half of the trials. At the end of each trial, an auditory response cue (either in a high tone or a low tone) informed observers about the location of the target. Observers then reported the orientation of the target using the keyboard. The neutral condition and the saccade condition were tested in separate blocks in a randomized order. In the saccade blocks, equal number of trials of the Toward and Away conditions were randomly interleaved. Observers were explicitly told that the direction of the saccadic cue was not informative regarding the target location.

There were three conditions to investigate the effect of covert endogenous attention: neutral, valid and invalid (**Figure 3A**). The procedures in these conditions were similar to the procedures of the neutral, saccade Toward and saccade Away conditions described above, except for the following changes. The SOA between the pre-cue and the test stimulus was 400 msec so that the effect of covert endogenous attention could take full effect (reviewed in Carrasco, 2011). For all the conditions, observers had to maintain fixation at the screen center throughout each trial; eye position was monitored throughout the trials. There were two types of blocks, one containing only the neutral condition, and the other containing both the valid and invalid conditions. In the latter, 70% of the trials were valid trials (with the pre-cue pointing toward the target) and the remaining 30% were invalid trials (with the pre-cue pointing toward the distractor). Observers were told that the pre-cue was informative regarding the target location, and that they should pay attention to the cued location.

The covert exogenous attention task was similar to the endogenous attention task, except the following modifications. The pre-cue was presented peripherally and transiently with a duration of 60 ms. The pre-cue was followed by an inter-stimulus interval (40 ms) before the test stimulus. This led to a 100-ms SOA between the pre-cue and the test stimulus, which yields the greatest effect for exogenous attention (reviewed in Carrasco 2011). In the valid condition, the pre-cue was a black line (0.5° length) presented above the target (1.8° above the center of the target), whereas in the invalid condition, the pre-cue was presented above the distractor. In the neutral condition, the pre-cue were black bars presented at the fixation, equivalent to the neutral cue in the saccade and the endogenous attention tasks. Similar to the endogenous attention task, there were two types of blocks, one containing only the neutral condition, and the other containing both the valid and invalid condition. In the latter, equal number of valid and the invalid trials were randomly interleaved. Observers were told that the pre-cue was uninformative regarding the target location. Eye position was monitored throughout the trials.

### Experiment 3

We repeated the presaccadic attention task in Experiment 2 with increased location uncertainty. Again, we instructed observers to saccade to the center of the placeholder in the saccade condition. There were still two placeholders on the screen, but now with five potential target locations within each placeholder. We considered the presaccadic attention task in Experiment 2 as a low-uncertainty condition and we tested two levels of location uncertainty here, named as medium-uncertainty (**Figure 5A**) and high-uncertainty (**Figure 5B**) conditions that differed in the area covered by the placeholders. In each trial, the target was presented at any one of the five locations in a placeholder with equal probability. Similarly, the distractor was presented at any one of the five locations in the other placeholder. The location of the test stimuli varied across trials. On average, every observer completed 7722 trials in 10 sessions on different days.

### Stimuli

The test stimuli were Gabor patches generated by a sinusoidal function (4 cycle per degree with its phase randomly determined in each trial) weighted by a Gaussian with a standard deviation of 0.5 degree. Test stimuli in 9 contrast levels (5, 10, 14, 20, 27, 37, 51, 70, 95%) were tested. Contrast levels were randomly selected on each trial, and were interleaved in each of the experiments. In Experiments 2 and 3, the target and the distractor in a trial were always presented with the same contrast level.

The orientation of the Gabors was titrated for each observer. The titration procedure was conducted using the procedure of the neutral condition of each experiment and task. Three types of tasks (presaccadic attention, endogenous attention and exogenous attention) in Experiment 2 were titrated separately. The medium-uncertainty and high-uncertainty condition in Experiment 3 were also titrated separately. Test stimuli were presented at the highest contrast level and an adaptive procedure (Watson and Pelli, 1983) was used to determine the amount of tilt (from vertical) at which observers’ performance in discriminating the orientation (left vs. right tilt) of the target was at about d′=2.

Two placeholders (centered at 8° left and right from the fixation) were presented on the screen throughout all the experiments. The placeholders were composed of four black dots forming a square with a width of 2.3° in Experiment 1 and 2. In Experiment 3, the placeholders had a width of 3.4° and 6.3° in the medium uncertainty and large uncertainty conditions, respectively.

In Experiments 1 and 2, the test stimuli were presented at 8° left or right from the fixation. In Experiment 3, the test stimuli were also presented 8° from fixation but could occur at any one of the five locations within the placeholder. The five locations were evenly spaced on an imaginary circular arc symmetric around the horizontal meridian. The imaginary arc had a radius of 8° and an angle of 18° and 44° in the medium- and the high-uncertainty conditions, respectively. Thus, the two outmost locations (the top and the bottom) had a distance of 2.5° and 6° in the medium and the high-uncertainty conditions, respectively (see **Figure 5A and 5B**, in which the locations are marked by dashed lines for illustration purposes). Observers were told to saccade to the center of the placeholder.

### Analysis

We fitted psychometric functions (*d*′ as a function stimulus contrast) with Naka-Rushton functions parameterized as *d′=d*_*max*_*C*^*n*^*/(C*^*n*^*+C*^*n*^_*50*_) where *C* is the contrast level, *d*_*max*_ is the asymptotic performance, *C*_*50*_ is the contrast level corresponding to half the asymptotic performance, and *n* determines the slope of the function. A change *d*_*max*_ indicates a response gain change and a change of *C*_*50*_ indicates a contrast gain change.

A bootstrapping procedure was used to test the statistical significance of the change of *d*_*max*_ and *C*_*50*_ across conditions. We first resampled (with replacement) the data to obtain a psychometric function (nine data points) for each observer and condition. We then computed group-averaged psychometric functions by averaging the data points across observers. Naka-Rushton functions were then fitted to the group-averaged psychometric functions. We subtracted the *d*_*max*_ (or *C*_*50*_) between valid and invalid conditions to obtain the difference score. This procedure was repeated 2000 times to generate the distribution of difference scores. The *p*-values were computed as the proportion of the difference scores that fell below or above zero.

### Eye Position

We monitored eye positions online. Trials with blinks or saccades (except for the response saccade required in the saccade condition) detected from the beginning of the trial to 200 ms after the offset of the test stimuli were aborted and repeated at the end of the block. For offline analysis, we smoothed the raw eye data with a Gaussian and computed smoothed eye velocity using the eye positions of five neighboring time points. Saccades were detected when the eye velocities exceeded the median velocity by 5 standard deviations for at least 8 ms (Engbert and Mergenthaler, 2006). Saccades separated by less than 10 ms were merged as a single saccade. For the saccade condition, we analyzed the trials in which observers’ first response saccade occurred between 70–400 ms after the cue onset, and landed within the cued location with an error smaller than 2.5° (from the center of the aperture) in Experiment 1 and 2, and smaller than 3.5° in Experiment 3.

### Models

#### Normalization model of attention (NMA)

We simulated population neural responses following Reynolds and Heeger’s Normalization Model of Attention (2009). We simulated a 2-dimensional array of neurons with orientation tiling the orientation space in steps of 1°, and receptive field centers ranging from −10° (degree visual angle) to 10° relative to the target location in steps of 0.5°. The responses of the neurons (*R*) were computed as

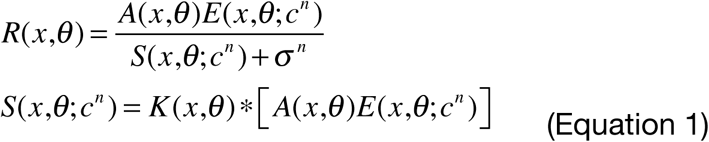

where *E* is the excitatory drive, *S* is the suppressive drive, *A* is the attentional gain factor, *c* is stimulus contrast, *n* is an exponent term and *σ* is a nonnegative suppression constant (**Figure 7A**). The excitatory drive *E* was determined by each neuron’s receptive field center *x* and preferred orientation *θ*. The spatial excitatory field of each simulated neuron was a Gaussian with 1° standard deviation and the orientation tuning was a circular Gaussian with 48° FWHM (full width at half maximum), approximating the spatial and orientation tunings reported for macaque primary visual cortex (Graf et al., 2011; Keliris et al., 2019). Suppressive drive *S* was computed as a suppression kernel *K* convolved with attentional-modulated excitatory drive. *K* is a 2-dimensional kernel in which the spatial extent was modeled as a Gaussian with a width four times that of the spatial excitatory field, emulating the broad suppressive surround of visual neurons. For simplicity, we set *K* to be uniform across orientations, meaning that at the same locations, the normalization pool was summed across all the neurons.

Attentional modulation was modeled by attentional gain factor (*A*) that multiplicatively modulates the excitatory drive before normalization. Attentional gain factors are uniform across orientation (**Figure 7A**), and the spatial extent of the attentional gain factor is modeled as a Gaussian,

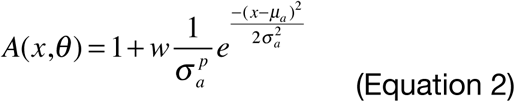

where 1 is the baseline of the attention gain factor, *μ*_*a*_ is the attended location (assumed to be centered at the central target location), *σ* _*a*_ determines the size (width) of the spatial extent of the attentional gain factor, and *p* is a parameter controlling the tradeoff between the spatial extent and the magnitude of attentional modulation. If *p=1*, spatial attention has a fixed volume, decreasing in magnitude when its spatial extent increase. If *p=0*, the magnitude of the attentional gain is independent of the spatial extent of attentional modulation. *w* is a weight that determines the strength of attentional modulation. We fitted this weight with two free parameters *w*_*n*_, *w*_*t*_, corresponding to the strength of attention in the neutral and the Toward (or the valid) condition. The weight of the Away (or the invalid) condition was fixed at zero.

Behavioral performance (*d*′) in a discrimination task is proportional to the neural responses given the assumption of an additive, independent, and identically distributed (IID) noise. An alternative model with Poisson noise and a maximum-likelihood decision rule would yield the same linkage between neural response and behavioral performance (Jazayeri and Movshon, 2006; Pestilli et al., 2009). Therefore, we fitted a free parameter, *σ* _*n*_, representing the magnitude of the noise, to relate the underlying neural responses to behavioral reports.

We computed probabilistic behavioral report using the simulated population neural response and signal detection theory. The available signal *s* for an observer is computed as the Euclidean norm of the subtraction between two vectors *s* = ||**r**_*R*_ – **r**_*L*_ ||, in which **r**_*R*_ represents the population neural response for a right-tilted target and **r**_*L*_ represents the population neural response for a left-tilted target. Based on signal detection theory, the observer’s performance is *d*’ = *s / σ* _*n*_, and the probability for the observer to correctly discriminate a right-tilted target is 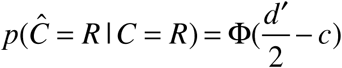, where *C* is the true category of the target (left-tilted *L* or right-tilted *R*), Ĉ is the observer’s response, *c* is a response bias (i.e., an idiosyncratic preference for reporting a particular category) and Φ is cumulative distribution function of the standard normal distribution.

The inputs to the model were based on the stimuli in the experiments. We simulated population neural responses (**r**_*R*_ and **r**_*L*_) for the right-tilted orientation and for the left-tilted orientation by setting the orientation of the input stimulus at 4° and −4° degree relative to vertical respectively. For the medium and high location uncertainty conditions, the neural responses for the outmost locations, intermediate locations and the central location were simulated separately by moving the location of the input stimulus based on our stimuli (**Figure 5A**).

Below, we consider alternative models. All the models share the same implementations as the NMA described above, but differ in how and when attention modulates neural responses.

#### Response gain model (RG)

The effect of attention is modeled as the response gain factor *γ* that scales the response of the neurons multiplicatively after normalization. The responses of the neurons (*R*) are computed as

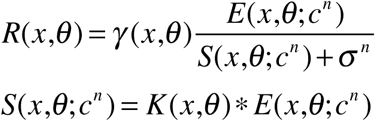

#### Contrast gain model (CG)

The effect of attention is modeled as the contrast gain factor *ϕ* that divisively modulate the suppression constant of the neuron. *ϕ* horizontally shifts the response functions of the neurons left (*ϕ* >1) or right (*ϕ* <1). The responses of the neurons (*R*) are computed as

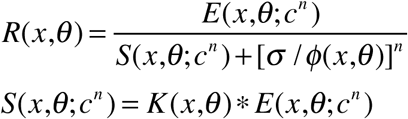

#### Input baseline model (IB)

Here attention is modeled as a change of baseline input *I*. The response of the neurons (*R*) are computed as

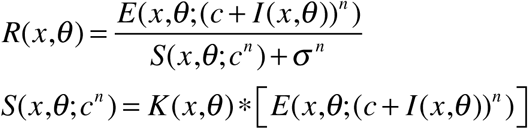

We found that an input baseline added after the exponent term *E*(*x,θ*; *c*^*n*^) + *I* (*x,θ*) to fit the data almost equally well as the input baseline added prior to exponentiation. Thus, we only report one version of this model.

#### Output baseline model (OB)

Attention is modeled as an additive term *O* after normalization. This is similar to the ‘increased baseline’ observed in previous human fMRI studies on covert spatial attention (Li et al., 2008).

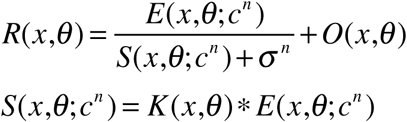

Response gain factor *γ*, contrast gain factor *ϕ* are all modeled using Equation 2 with the same free parameters *p* and *σ* _*a*_. Baseline input *I* and output baseline *O* also follow Equation 2, except that they have a baseline at 0, instead of 1. Note that all these variables (*γ, ϕ, I, O*) can be conceptualized as attentional gain factors. Here, we use different names and symbols to distinguish them from the terms used in NMA.

#### Model variants

In addition to different models of attentional modulation (NMA, RG, CG, IB and OB), we varied other factors in the model: (1) The response bias *c* was either fitted as a free parameter or fixed at zero (no bias). (2) The trade-off between the size of attentional field and the magnitude of attention was either allowed to be a free parameter (*p* in Equation 2), or fixed at 0 (no trade-off).

We conducted factorial model comparison using all the factors above leading to 20 models in total: 5 types of attentional modulations (NMA, RG, CG, IB, OB) X bias (B) or no bias (NB) allowed X trade-off (T) or no trade-off (NT) allowed (**Supplementary Table 1**). This procedure ensures that the winning model does not depend on the setting of the models we choose a priori.

#### Model Fitting

We denote the free parameters collectively by *θ* (free parameters of each model are listed in **Supplementary Table 1**). We fit each model to individual-subject data by maximizing the log likelihood of *θ*, log *L*(*θ*) = log *p*(data | *θ*). We assume that the trials are conditionally independent. The log likelihood is 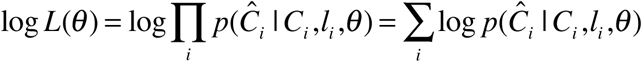, where *l*_*i*_ is the location of the target, *C*_*i*_ the true category (right-tilt or left-tilt) of the target and *Ĉ*_*i*_ represent observers’ response on the *i*th trial respectively. We optimized the parameters for each individual using Bayesian Adaptive Direct Search (Acerbi and Ma, 2017). To select the best model, we computed AIC (Akaike information criterion) for each observer and model. The best-fit model is the model with the lowest group-summed AIC. We reported group-summed ΔAIC (the AIC of the models minus the AIC of the best-fit model) and its confidence interval estimated by bootstrapping.

#### Simulation prior to Experiment 3

Before running Experiment 3, we ran a simulation to investigate the predictions of NMA. In this simulation, the model parameters were handpicked. We scaled the size of attention field with the size of the placeholders, and we set the weight of the attentional modulation so that the ratio between the asymptotes of the simulated neural response in the Toward and the Away condition (under the low location uncertainty) was similar to the ratio of *d*_*max*_ between the Toward and Away conditions in the data.

## Supplementary Materials

**Supplementary Figure 1.**
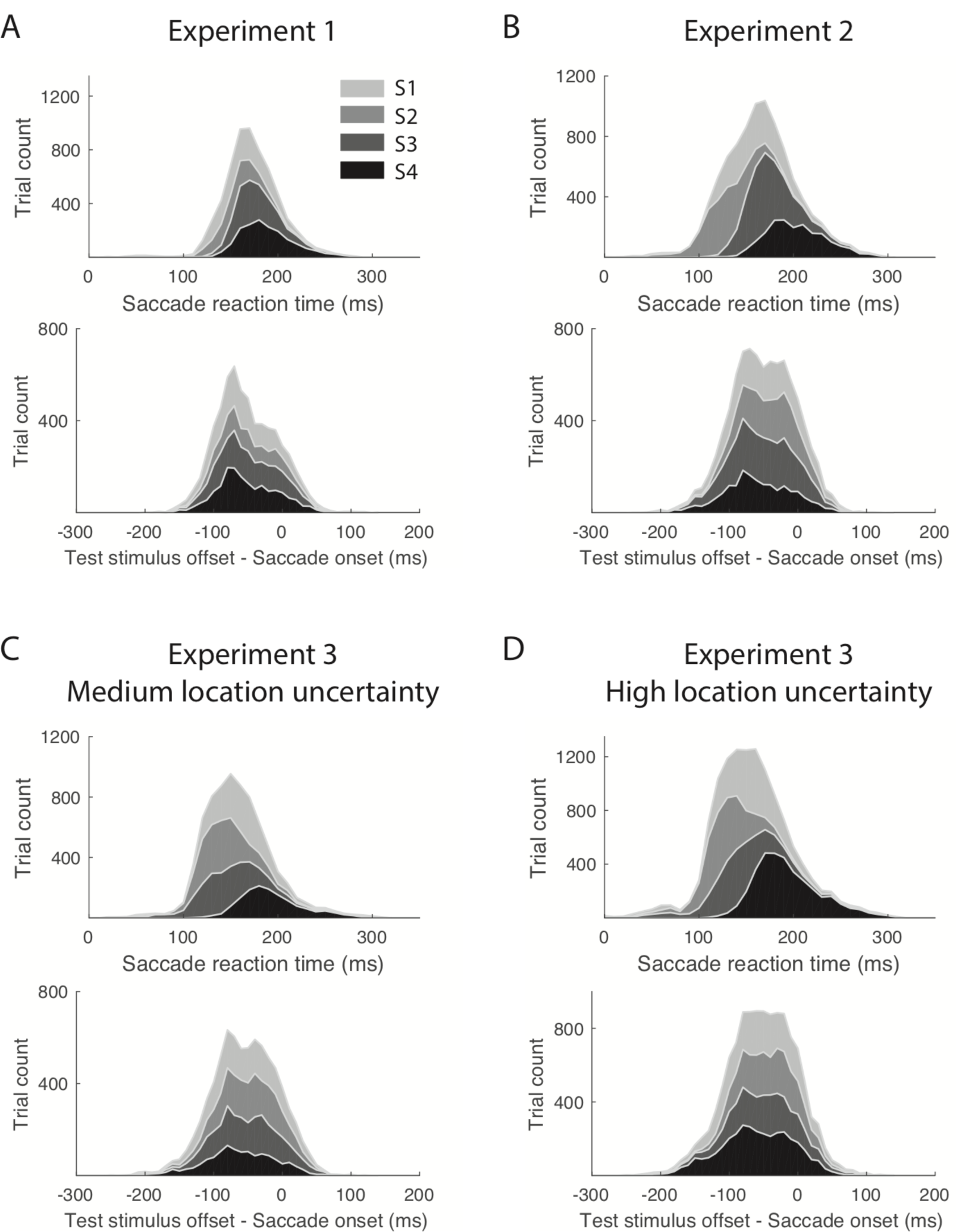
Distribution of saccade reaction time (saccade onset minus saccade cue onset) and latency (test stimulus offset minus saccade onset). **(A)** Experiment 1. **(B)** Experiment 2. **(C)** Experiment 3, Medium location uncertainty condition. **(D)** Experiment 3, High location uncertainty condition.

**Supplementary Figure 2.**
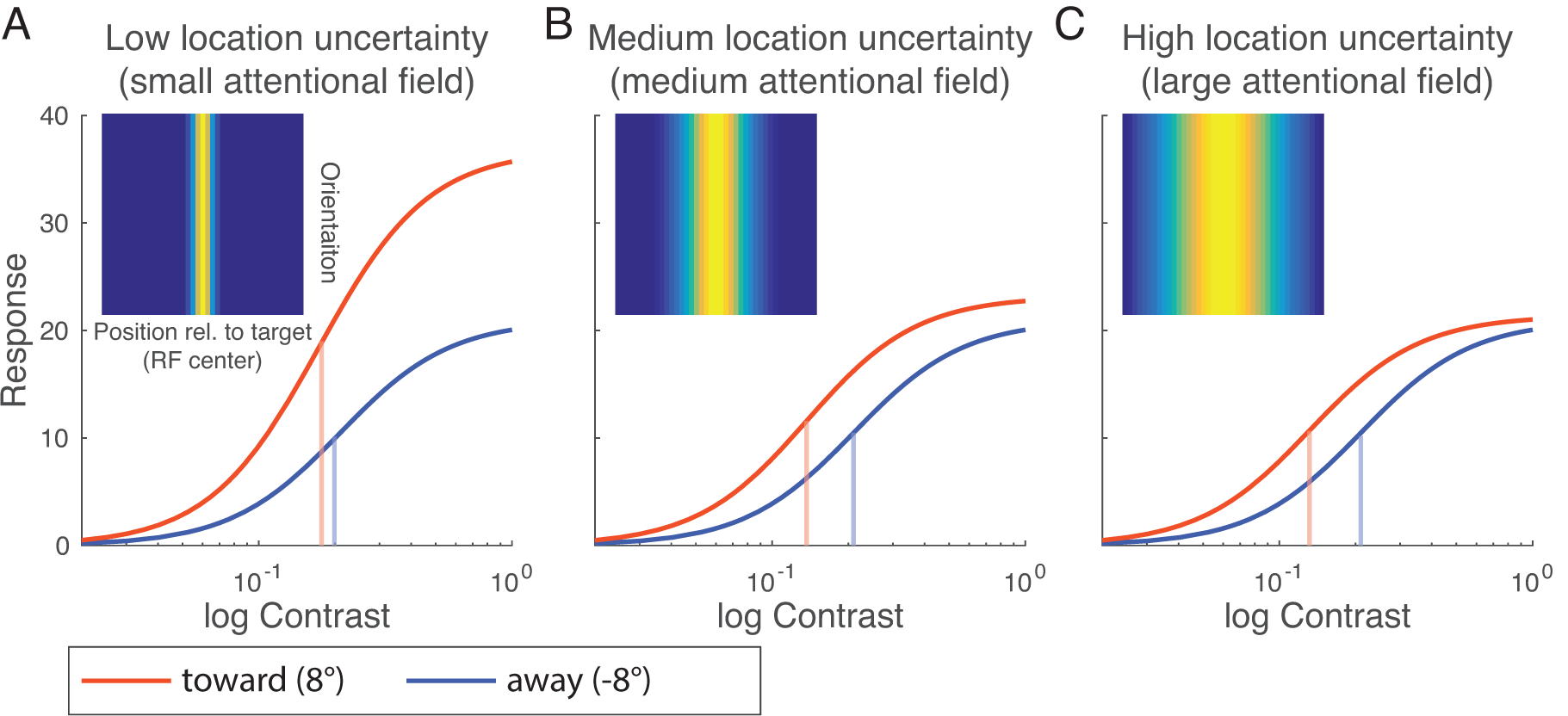
Simulations. Neural responses were simulated based on the Normalization Model of Attention (Reynolds and Heeger, 2009; See NMA in Models in Methods). (**A**) Simulated neural response as a function of stimulus contrast in the low-uncertainty condition. The vertical lines indicate the semi-saturation contrast of the contrast response functions. The legend is the attentional gain factor in this condition. The inset is the attentional gain factors used for simulating the Toward condition. The attentional gain factors were set at the baseline (=1, equivalent to the dark blue color in the inset) across the entire neural population for the away condition. (**B**) Medium location uncertainty condition. (**C**) High location uncertainty condition.

**Supplementary Figure 3.**
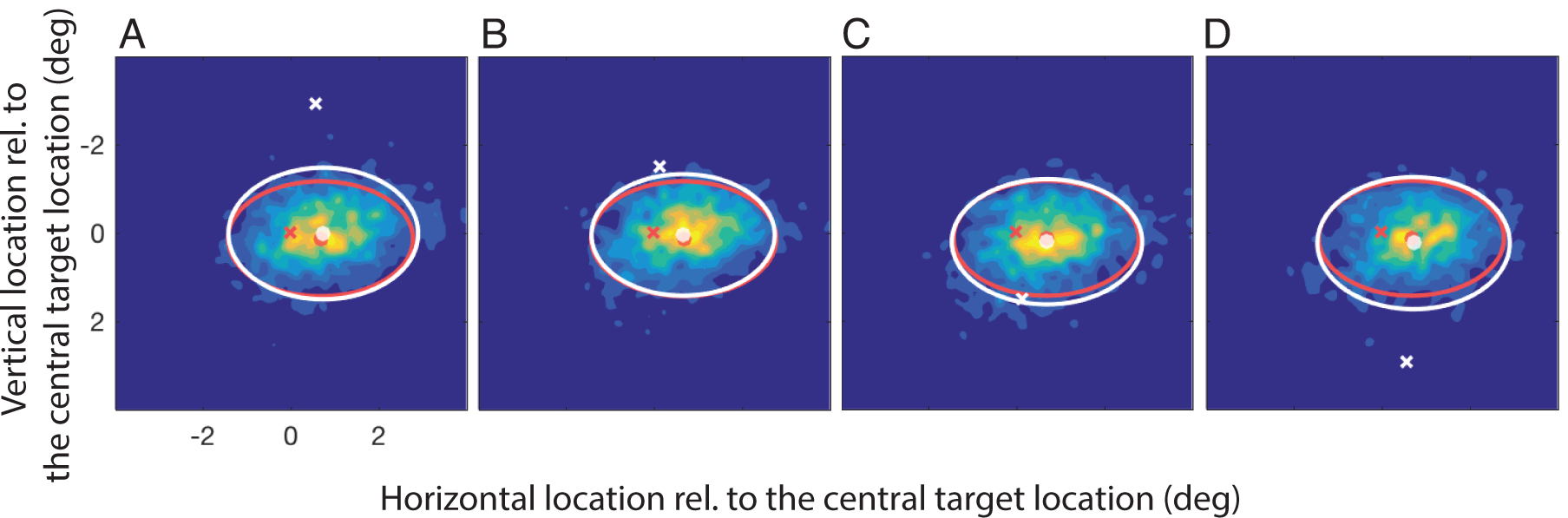
Distributions of saccade landing positions in the high-uncertainty condition in Experiment 3. We collapse leftward and rightward saccades by flipping the sign of the horizontal location of the saccade landing points of all the rightward saccades. Distributions of saccade landing points were first computed for each observer and then averaged across all observers. In all the figures, we compared the distribution of the saccade landing positions when the target was presented at the off-center locations with the distribution when the target was presented at the central location. In all the figures, the red cross is the location of the central target, and the red ellipse and the red dot represent the distribution of saccade landing positions when the target was presented at the central location. (**A**) The background color represents the distribution of saccade landing positions when the target was presented at the topmost position (denoted by the white cross). This distribution is further illustrated by the white ellipse: The center of the white ellipse is the mean of the distribution (denoted by the white dot), and the axes of the white ellipse represent ±standard deviation along horizontal and vertical directions. (**B**-**D**) In the same format as (**A**), the distributions of saccade landing points when the target was presented at the other three off-center locations were illustrated by the background color and all the white symbols.

**Supplementary Figure 4.**
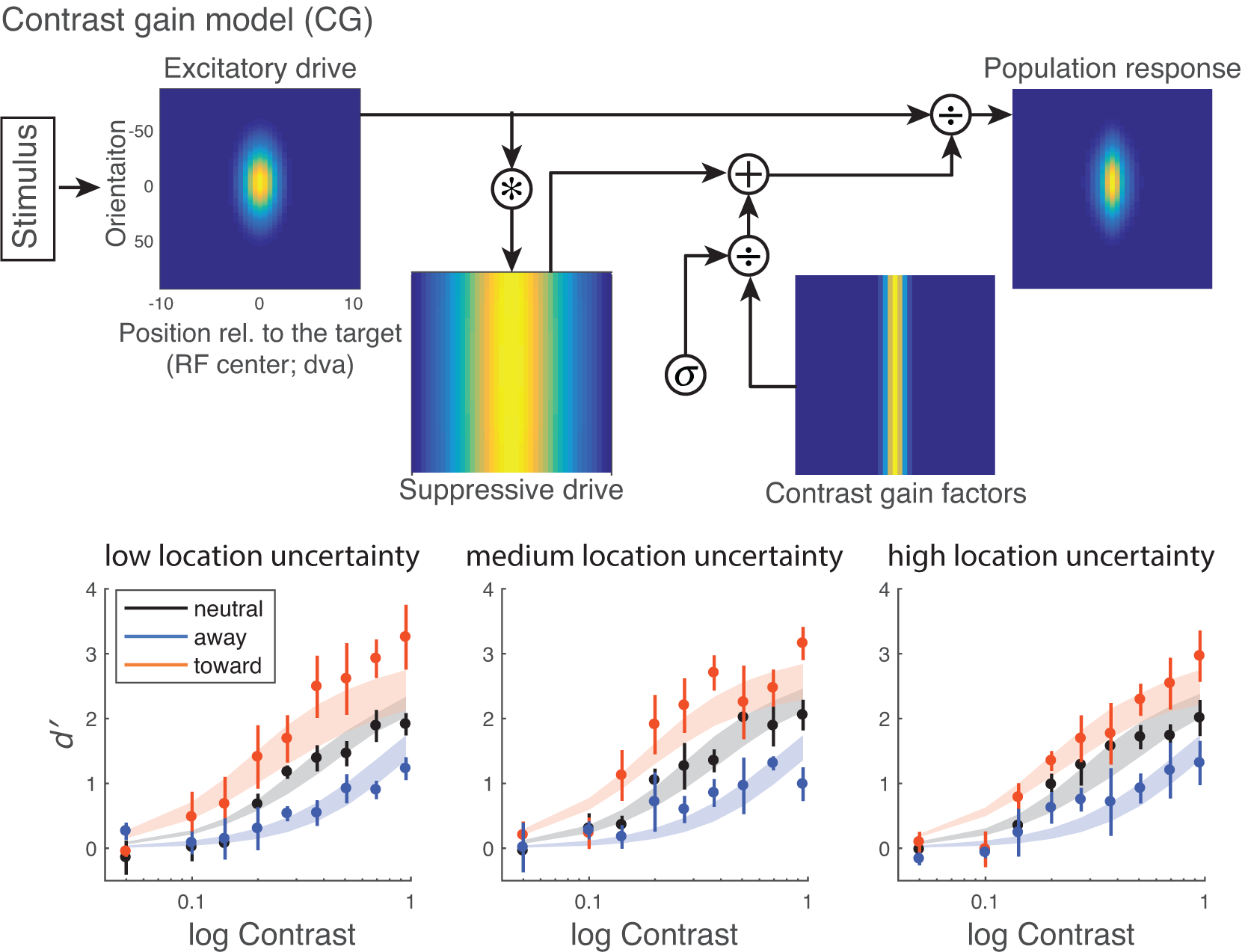
Model and model fit. Top row: Contrast gain model. Attention modulates the suppression constant (*σ*) of the neurons divisively. Note that the suppression constant is present in all the models, but for simplicity, it is not shown in the figures of other models. Bottom row: Model fit. Data points and error bars represent group-averaged data and ±1 s.e.m. The model was fitted to individual subjects, and the shading areas represent the averaged of the fit across participants (mean ±1 s.e.m).

**Supplementary Figure 5.**
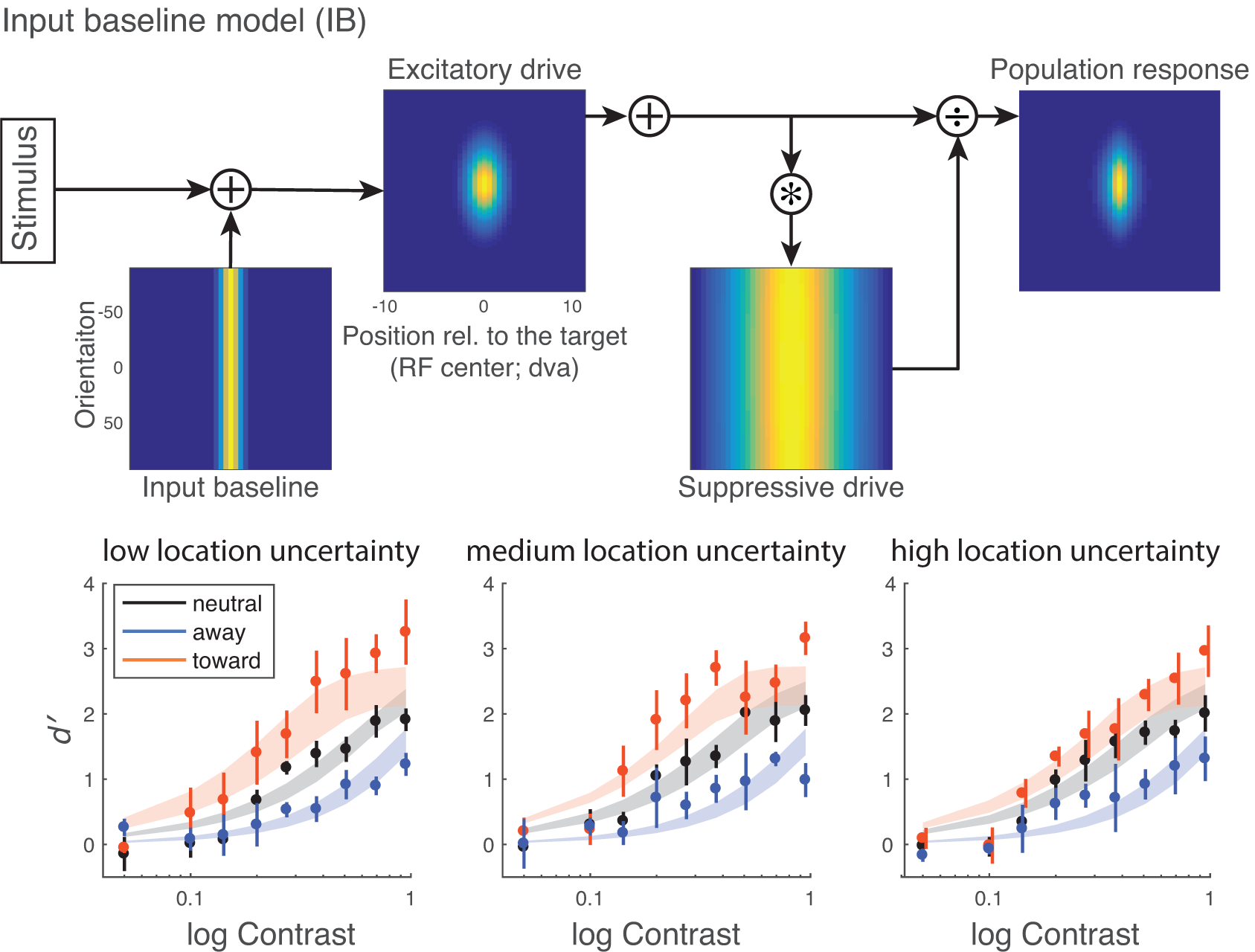
Model and model fit. Top row: Input baseline model. Attentional modulation modeled as an additive term at the input baseline of the neuron. Bottom row: Model fit. Data points and error bars represent group-averaged data and ±1 s.e.m. The model was fitted to individual subjects, and the shading areas represent the averaged of the fit across participants (mean ±1 s.e.m).

**Supplementary Figure 6.**
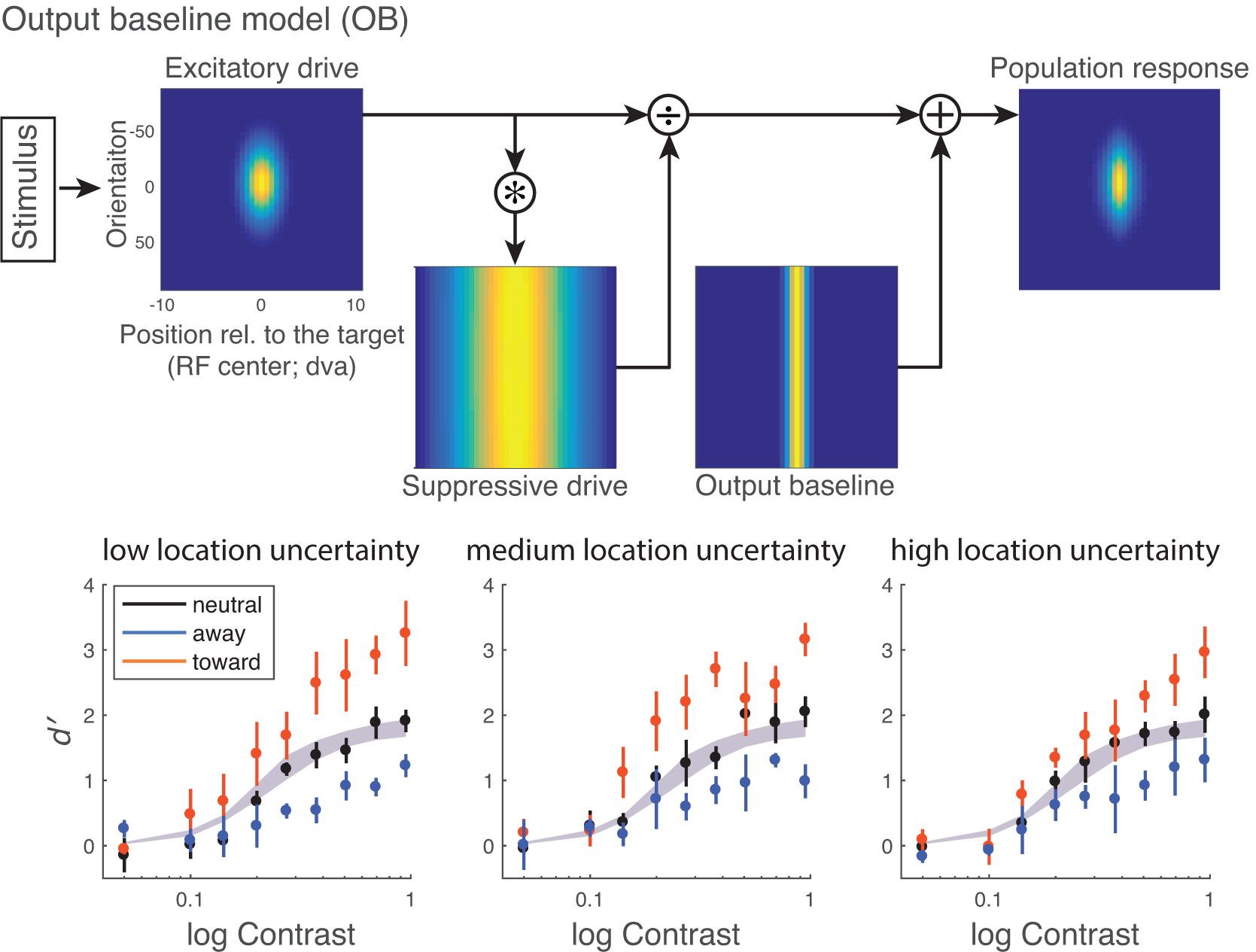
Model and model fit. Top row: Output baseline model. Attention is modeled as an additive term after the normalization. Bottom row: Model fit. Data points and error bars represent group-averaged data and ±1 s.e.m. The model was fitted to individual subjects, and the shading areas represent the averaged of the fit across participants (mean ±1 s.e.m).

**Supplementary Figure 7.**
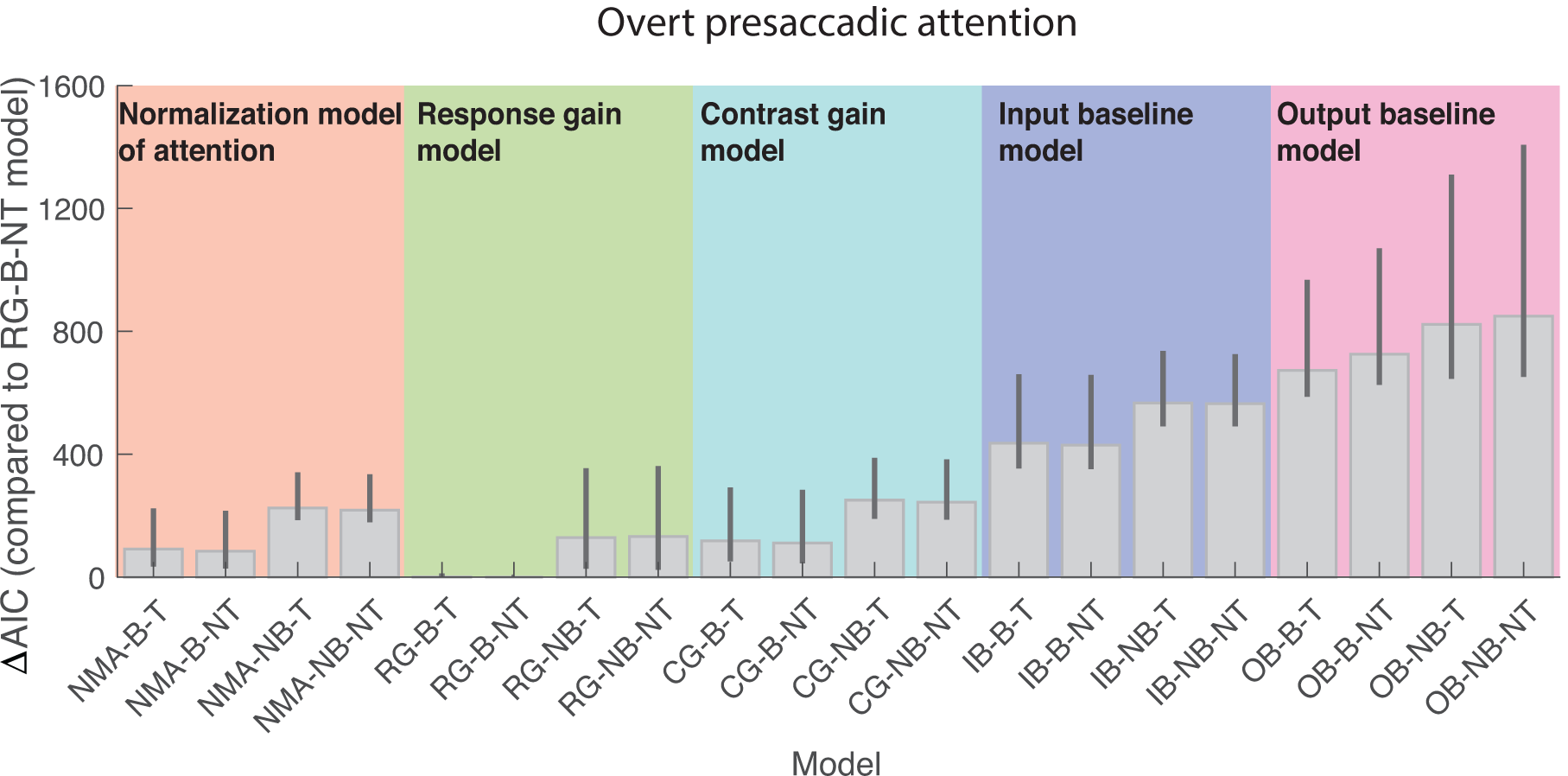
Factorial model comparison for overt presaccadic attention using AIC. ΔAIC is the AIC of each model minus the AIC of the best-fit model (RG-B-NT). The bars represent ΔAIC summed across participants. The error bars represent 95% bootstrapped confidence interval. Names of the model are denoted as the attentional modulation paired with different implementations of other factors in the model separated by hyphens (-). NMA: normalization model of attention. RG: response gain; CG: contrast gain; IB: input baseline; OB: output baseline; B: response bias allowed; NB: no response bias allowed; T: trade-off between attention field size and the strength of attention allowed; NT: no trade-off between attention field size and the strength of attention allowed.

**Supplementary Figure 8.**
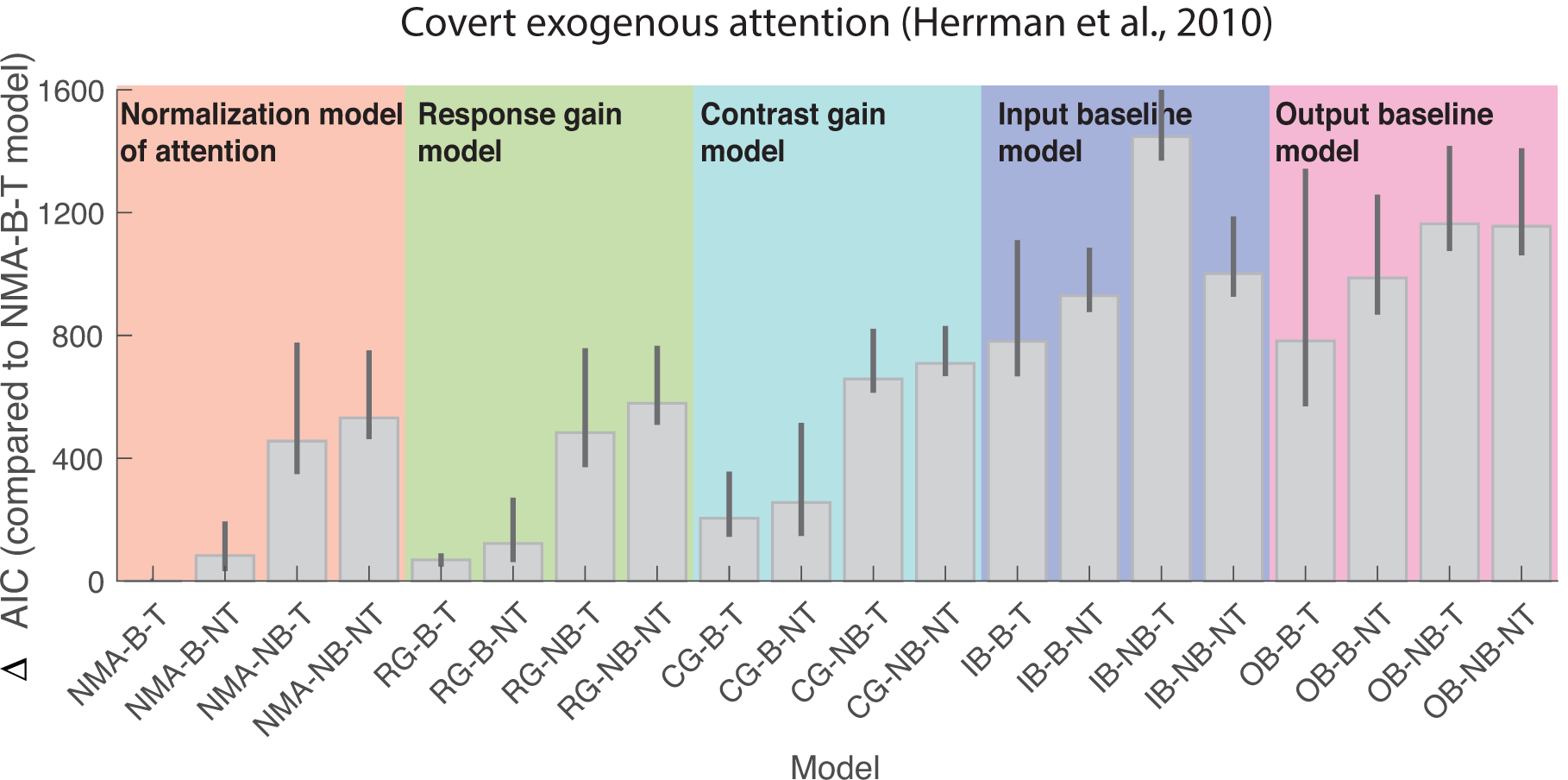
Factorial model comparison for covert exogenous attention using AIC. ΔAIC is the AIC of each model minus the AIC of the best-fit model (NMA-B-T). The bars represent ΔAIC summed across participants. The error bars represent 95% bootstrapped confidence interval. Names of the model are denoted as the attentional modulation paired with different implementations of other factors in the model separated by hyphens (-). NMA: normalization model of attention. RG: response gain; CG: contrast gain; IB: input baseline; OB: output baseline; B: response bias allowed; NB: no response bias allowed; T: trade-off between attention field size and the strength of attention allowed; NT: no trade-off between attention field size and the strength of attention allowed.

**Supplementary Figure 9.**
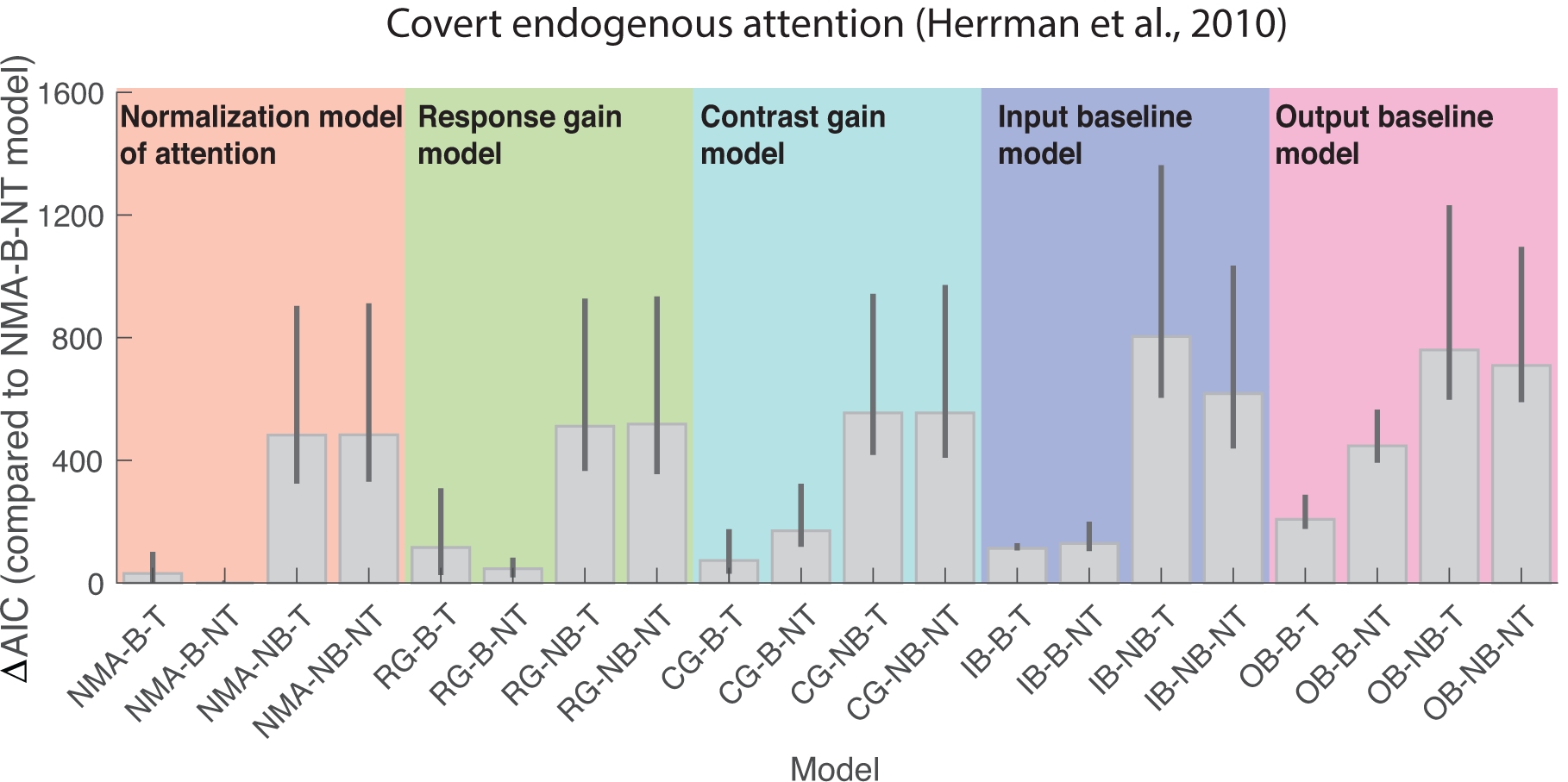
Factorial model comparison for covert endogenous attention using AIC. ΔAIC is the AIC of each model minus the AIC of the best-fit model (NMA-B-NT). The bars represent ΔAIC summed across participants. The error bars represent 95% bootstrapped confidence interval. Names of the model are denoted as the attentional modulation paired with different implementations of other factors in the model separated by hyphens (-). NMA: normalization model of attention. RG: response gain; CG: contrast gain; IB: input baseline; OB: output baseline; B: response bias allowed; NB: no response bias allowed; T: trade-off between attention field size and the strength of attention allowed; NT: no trade-off between attention field size and the strength of attention allowed.

**Supplementary Table 1.** Free parameters of the model.

| Free parameters | Descriptions |
| --- | --- |
| $\sigma$ | suppression constant of the neurons |
| $n$ | exponent term of the neurons |
| $\sigma_n$ | the magnitude of neural noise |
| $w_n$ | the strength of attention in the neutral condition |
| $w_t$ | the strength of attention in the Toward condition |
| $\sigma_{aL}$ | the width of attention field in the low uncertainty condition |
| $\sigma_{aM}$ | the width of attention field in the medium uncertainty condition |
| $\sigma_{aH}$ | the width of attention field in the high uncertainty condition |
| $c$ | response bias |
| $p$ | the trade-off between the strength of attentional modulation and the size of attentional field |
| All the models contain free parameters $\sigma, n, \sigma_n, w_n, w_t, \sigma_{aL}, \sigma_{aM}, \sigma_{aH}$. The other two parameters, $c$ and $p$, were either set as a free parameter or fixed at 0 depending whether the model allows a response bias, a trade-off term or both.<br>The strength of attention in the Away (or invalid) condition was fixed at zero as the baseline. The width of attention field was constrained as $\sigma_{aL} \leq \sigma_{aM} \leq \sigma_{aH}$. Herrmann et al. (2010) only tested two levels of location uncertainty, and thus only two free parameters ($\sigma_{aL}, \sigma_{aH}$) were used for the width of attention field. |  |

